# Cell State-Specific Cytoplasmic Material Properties Control Spindle Architecture and Scaling

**DOI:** 10.1101/2024.07.22.604615

**Authors:** Tobias Kletter, Omar Muñoz, Sebastian Reusch, Abin Biswas, Aliaksandr Halavatyi, Beate Neumann, Benno Kuropka, Vasily Zaburdaev, Simone Reber

## Abstract

Mitotic spindles are dynamically intertwined with the cytoplasm they assemble in. How the physicochemical properties of the cytoplasm affect spindle architecture and size remains largely unknown. Using quantitative biochemistry in combination with adaptive feedback microscopy, we investigated mitotic cell and spindle morphology during neural differentiation of embryonic stem cells. While tubulin biochemistry and microtubule dynamics remained unchanged, spindles changed their scaling behaviour: in differentiating cells, spindles were significantly smaller than those in equally-sized undifferentiated stem cells. Integrating quantitative phase imaging, biophysical perturbations and theory, we found that as cells differentiated, their cytoplasm became more dilute. The concomitant decrease in free tubulin activated CPAP (centrosomal P4.1-associated protein) to enhance the centrosomal nucleation capacity. As a consequence, in differentiating cells, microtubule mass shifted towards spindle poles at the expense of the spindle bulk, explaining the differentiation-associated switch in spindle architecture. This study shows that cell state-specific cytoplasmic density tunes mitotic spindle architecture. Thus, we reveal physical properties of the cytoplasm as a major determinant in organelle size control.

## INTRODUCTION

The mitotic spindle is a prime example to study subcellular organising principles and link physical laws to organelle function ^1^. While spindles showcase a staggering morphological diversity, their primary function remains a mechanical one: to precisely partition the genetic material into two daughter cells ^2^. Generally, spindles show two scaling regimes adapting to changes in cell size: in small cells spindle size scales linearly with cell size. In large cells, the size of the spindle becomes decoupled from cell size and reaches an upper limit. Such key observations of spindle scaling mostly stem from fast reductive divisions in early mouse, frog, fish, and worm embryos ^3–10^. However, later in development, cellular differentiation poses a similar challenge: differentiating cells change size, morphology, and function and therefore must reorganise and adapt their internal architecture. Changing spindle architecture while maintaining spindle function poses a structural and mechanical challenge to the cell and may require adaptation of the organising principles ^11^. Whether and how mitotic spindles adjust their morphology in differentiating cells remains largely unknown.

While spindles can function in a variety of cells across orders of magnitude of cell sizes, they all assemble mainly from one building block: αβ-tubulin. Tubulin assembles into dynamic microtubules, which self-organise with the help of microtubule-associated proteins (MAPs) and motors into a spindle with a steady-state length. How can more or less identical building blocks assemble spindles with varying sizes and architectures? Our current understanding is that small spindles modulate microtubule dynamics to adjust spindle size while large spindles require additional microtubule nucleation ^9,12,13^. Spindle microtubule nucleation and dynamics can be modulated in many ways, by changes in tubulin biochemistry ^14^, activity of MAPs and motors ^8,15–17^, by the presence of a limiting component ^18,19^ or changes in the biochemical composition of the cytoplasm ^6,9,20^. In the context of differentiation, adjustments to spindle architecture have been described ^21,22^. However, whether these architectural adjustments occur via modified microtubule dynamics or spatial regulation of microtubule nucleation is currently unknown. We therefore lack a systematic understanding how microscopic processes collectively give rise to a mesoscale spindle that is adaptive to the differentiating cellular context.

It is increasingly appreciated that spindles are dynamically intertwined with the cytoplasm they assemble in. However, mechanisms of how cytoplasmic properties affect spindle assembly and function are only starting to emerge. For example, it has been shown that cytoplasmic viscosity affects the rates of microtubule polymerisation and depolymerisation ^23^, spindle function ^24^, and spindle positioning ^25,26^. The mixing of the nucleoplasm and cytoplasm upon mitotic entry affects free tubulin concentration globally and consequently microtubule dynamics ^27^. At the same time, local enrichment of tubulin by organelle-exclusion zones ^28,29^ or at centrosomes ^30,31^ spatially supports microtubule nucleation and growth. In addition, the mitotic cytoplasm softens ^32^ and is fluidised by microtubule polymerisation dynamics ^33^. Both changes are thought to maintain diffusivity through the heterogenous metaphase cytoplasm. Although differentiation-mediated changes in cytoplasmic properties have been observed ^34–40^, we do not know how these changes directly affect spindle assembly and scaling.

Here, we use a cell culture model of neural differentiation to study changes in spindle architecture while cells change size, morphology, and function. Studying neural spindle scaling was motivated by the observation that neurodevelopment is particularly susceptible to mutations in spindle genes, and by pathologic evidence that links spindle morphology and mitotic susceptibility ^22,41–43^. To do so, we established a non-invasive, long-term image acquisition and analysis workflow that allowed us to image more than 4000 cells at single-cell resolution throughout neural differentiation. We find that spindle size subscales with cell volume: in early-differentiated cells spindles are 24% smaller when compared to spindles in equally-sized undifferentiated stem cells, but still follow a similar functional dependence on cell size. Despite this difference in size, tubulin biochemistry and microtubule dynamics remain unchanged. We show that cytoplasmic dilution and a drop in tubulin concentration increase centrosomal nucleation capacity. This is dependent on CPAP (centrosomal P4.1-associated protein), a centrosomal regulator ^44–46^. As a consequence, microtubule growth is redistributed to the spindle poles, resulting in a shift in spindle architecture. In agreement with this, differentiation-independent cytoplasmic dilution or the specific inhibition of the CPAP-tubulin interaction in undifferentiated ESCs phenocopied spindle architecture and size characteristic of early-differentiated cells. Our data are consistent with a theory in which microtubule number scales with cell volume and the inhibition of a centrosomal regulator determines the distribution between astral and bulk microtubules. To our knowledge, this is the first study to link cell state-specific cytoplasmic material properties to spindle architecture and size. More generally, this work shows how local intracellular environments can exert control over organelle morphology and scaling.

## RESULTS

### Spindle size subscales with cell volume upon differentiation

Mechanistic insights into spindle scaling stem primarily from fast, reductive cell divisions during early animal development ^3,4,8,9,13^. It thus remains largely unknown whether differentiating cells tune spindle size relative to cell size and if so, whether the observed physicochemical mechanisms apply to this cellular context. This question is particularly relevant in neurally differentiating cells where spindle defects have documented pathological implications ^41^. An experimentally accessible model is the differentiation of neurons from mouse embryonic stem cells (ESCs) in adherent cell culture ^47^. To quantitatively study spindle scaling, we differentiated ESCs that stably expressed Tubulin::GFP towards the neural lineage (Fig. 1a). This allowed us to image cells from pluripotency until terminal differentiation at single-cell resolution on a single substrate and in large numbers. We found that within 48 - 72 hours of neural induction, the differentiating cells downregulated the expression of the pluripotency marker OCT-4 and started expressing nestin (Fig. 1a-d) and PAX6 (Suppl. Fig. 1a), consistent with previous reports ^47,48^. Importantly, we observed the formation of neural rosettes (Suppl. Movie 1), which showed hallmarks of neuroepithelial physiology in terms of marker expression, cell polarity, and interkinetic nuclear migration^49,50^ (Fig. 1a, Suppl. Fig. 1b-g, Suppl. Movie 2). Thus, we concluded that this system represented a suitable model to study spindle morphology in a differentiation context.

**Figure 1:**
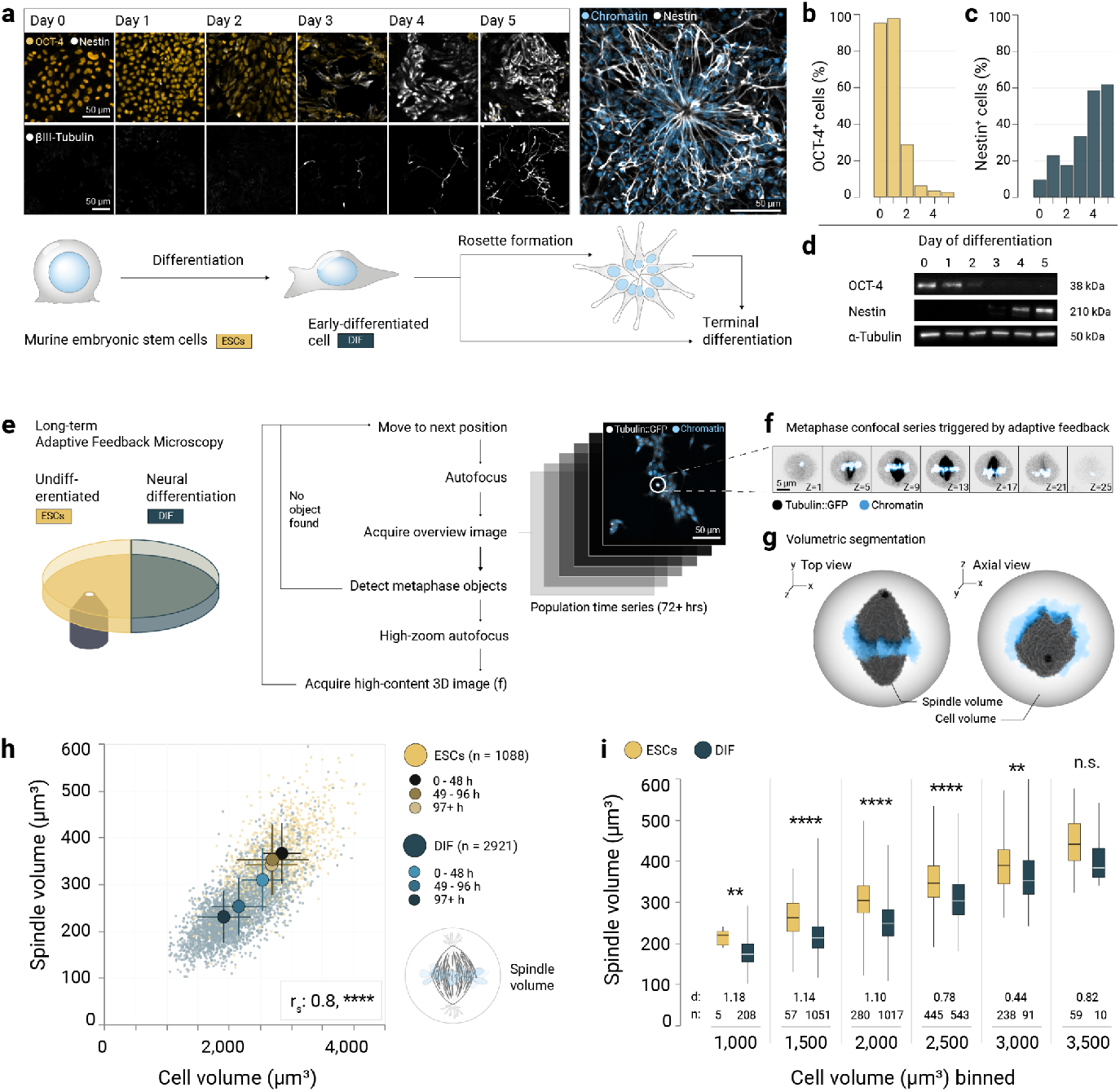
Spindle size subscales with cell volume upon differentiation. a) Immunofluorescence stainings of mouse embryonic stem cells (ESCs) driven towards neural fates in adherent monolayer. Left: every 24 hours, a replicate culture was stained for the pluripotency marker OCT-4 ^86^ (yellow, top row) together with the neural progenitor and rosette marker nestin (grey, top row), or the neuronal marker βIII-tubulin (grey, bottom row). Right: neural rosette after 6 days of differentiation stained with Hoechst (blue) and antibodies against nestin (grey). All images are maximum projections of confocal sections. Scale bars: 50 µm. Bottom: Illustration showing the differentiation of ESCs towards neural progenitor identities. b) Percentage of OCT-4-positive cells in the total cell population from a series of immunofluorescence stainings (as in a), covering the first 5 days of neural differentiation. c) Percentage of nestin-positive cells in the total cell population from a series of immunofluorescence stainings (as in a), covering the first 5 days of neural differentiation. d) Immunoblots probing for differentiation marker expression on a series of cell lysates, covering the first 5 days of neural differentiation. α-Tubulin as loading control. e) Automated microscopy experimental setup: tubulin::GFP-expressing ESCs were either kept undifferentiated in pluripotency-promoting conditions (“ESCs”) or driven towards neural differentiation (“DIF”). Fully automated adaptive feedback microscopy pipeline for live-cell imaging with high-content confocal series of metaphase cells expressing tubulin::GFP and chromatin stained with SiR-DNA. Scale bar: 50 µm. e) Confocal raw high resolution data of a metaphase cell captured using the adaptive feedback protocol as described above. Tubulin::GFP (inverted grey) and SiR-DNA (blue). Scale bar: 5 µm. f) 3D-rendered metaphase spindles (grey) after automated volumetric segmentation and morphometry using the *FIJI* plugin *Spindle3D* ^51^ and the pixel-classification tool *Ilastik* ^52^. Chromosomes are shown in blue. The cell volume is illustrated as a cartoon for clarity. g) Spindle volumes scale with cell volumes during differentiation. Each data point represents an individual cell (ESCs n = 1088 (yellow) and differentiation n = 2921 (blue)). Big circles represent the mean of each differentiation time bin. Error bars show the standard deviations. Data from N = 9 experiments. r_s_: Spearman’s correlation, ****: p < 0.0001. h) In equally-sized cells, spindle volume subscales in differentiating cells (blue) when compared to undifferentiated ESCs (yellow). Cell volume binning: 500 µm^3^. The white lines inside the boxes denote the medians, the boxes show the interquartile ranges, whiskers show the extrema. d: Cohen’s d (effect size), n: sample sizes in each bin. Welch’s t-test per bin, ****: p < 0.0001, **: p < 0.01, n.s.: not significant.

Using this model system, we designed an experimental and analytical methodology to track individual cell families through differentiation while simultaneously detecting and quantifying spindle morphology. Firstly, to overcome the challenge of generating sufficient quantitative three-dimensional (3D) imaging data of metaphase cells with minimal phototoxicity, we developed an adaptive feedback microscopy pipeline for live-cell imaging (Fig. 1e, Methods). Throughout the differentiation process, the fully autonomous microscope located metaphase cells within the larger population and launched high-content recordings exclusively of cells of interest (Fig. 1f). Next, using *Spindle3D* ^51^ and *Ilastik* ^52^, we extracted 3D morphometric parameters including spindle volume, cell volume, and cell surface area from 1,088 undifferentiated and 2,921 differentiating mitotic cells (Fig. 1g, h). Within the differentiating population, the average volume of mitotic cells decreased by 30% when compared to undifferentiated ESCs (V_ESC_ = 2719 ± 567 µm^3^, V_DIF_ = 1942 ± 424 µm^3^, Fig. 1h). This was independent of cell confluency and medium conditions (Suppl. Fig. 2a, b). Spindle volume decreased and scaled with cell volume (r_s_ = 0.8, Fig. 1h). However, spindles in the early-differentiating cells (up to 5 days of differentiation) occupied significantly less cell volume than spindles in undifferentiated ESCs (Suppl. Fig. 2c). This became even more apparent when we binned cell volumes: in cells with comparable cell volume, the median spindle volumes were up to 24% smaller in cells undergoing differentiation (Fig. 1i). This subscaling behaviour was independent of cell geometry or microtubule density within the spindle (Suppl. Fig. 2d-f). Thus, spindle size is not specified solely by cell volume. Taken together, we established a non-invasive, long-term image acquisition and analysis workflow, which allowed us to quantify cell and spindle morphologies over many generations to show that spindle volume subscales in early-differentiating cells. This meant that cell volume cannot be the only determinant of spindle size. Rather, our findings implied a cell state-specific spindle scaling mechanism that was responsive to the biochemical or physical changes imposed by cellular differentiation.

**Figure 2:**
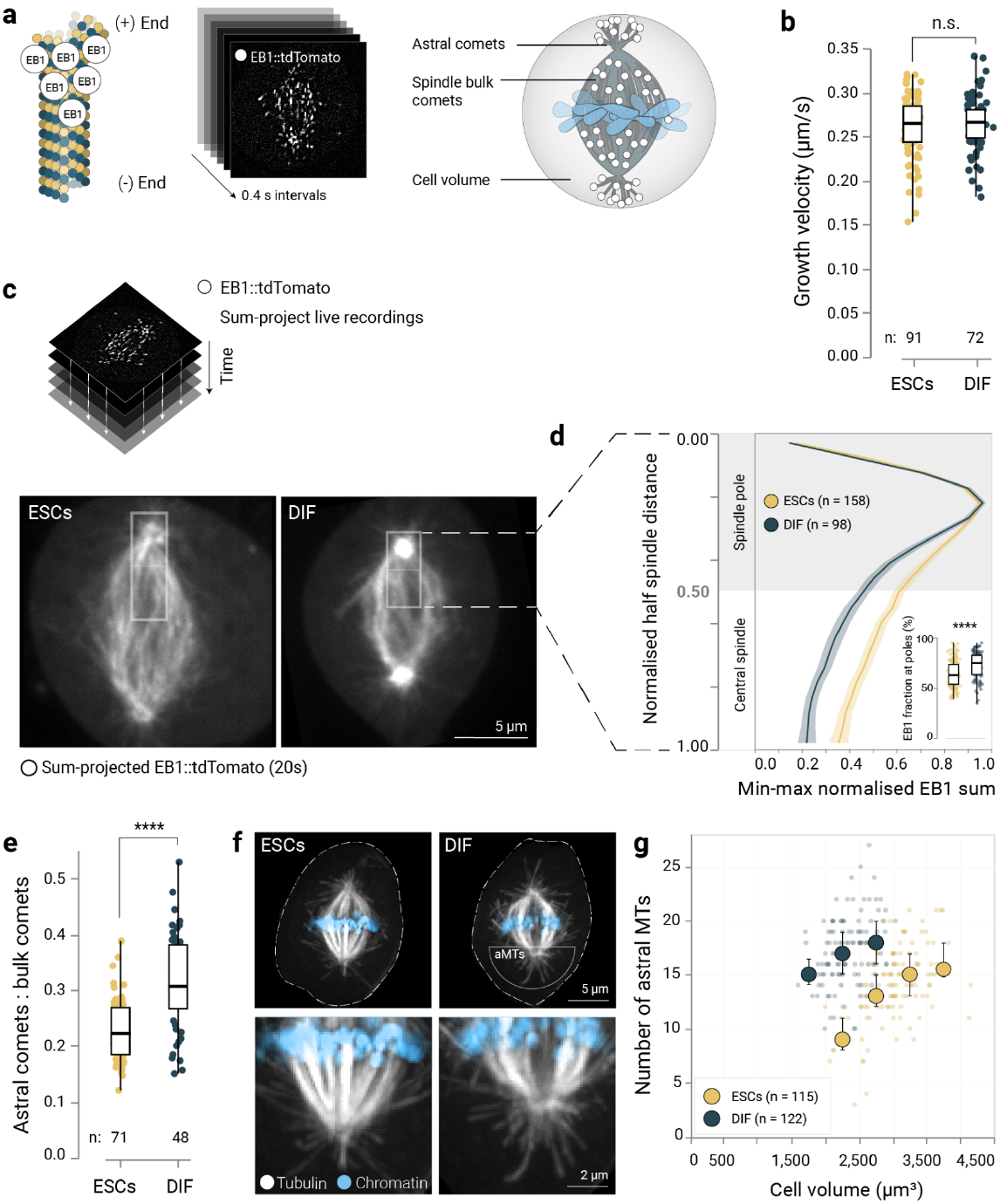
Spindles switch to pronounced astral architectures in early-differentiated cells. a) Mitotic cells transiently expressing EB1::tdTomato (a microtubule plus-end marker) were imaged live in 0.4 second intervals (left and center) to determine growth speed and distribution of growing microtubules before and after differentiation (right). b) The average growth speed of EB1::tdTomato-labelled microtubules (total comets). Data points show individual cells (ESCs n = 91, differentiation (48 h) n = 72 from N = 6 experiments). The boxes show the interquartile ranges, the black lines inside the boxes denote the medians, whiskers show the extrema. Welch’s t-test, n.s.: not significant. c) Top: EB1::tdTomato movies (20 s) were sum-projected. Bottom: Half-spindle intensity profiles were drawn spanning the distance between the spindle equator and the border between the spindle poles and the cytoplasm. d) EB1 concentration increases on spindle poles relative to spindle bulk in early differentiated cells. Intensity profile plots (see c) of the sum-projected EB1::tdTomato movies of ESCs and early differentiated cells were min-max normalised and pooled from N = 6 experiments. The lines show the means and the shaded areas show the 95% confidence intervals. Inset: the percentage of summed up EB1::tdTomato signal that is at the spindle pole (normalised distance < 0.5). Welch’s t-test, ****: p < 0.0001. e) The ratio of astral and spindle bulk EB1 comets changes during differentiation. Data points show the ratio in individual cells (ESCs n = 71, differentiation n = 48 from N = 6 experiments). The boxes show the interquartile ranges, the black lines inside the boxes denote the medians, whiskers show the extrema. Welch’s t-test, ****: p < 0.0001. f) Max-projected micrographs showing mitotic cells fixed and with immunostained microtubules (grey). Chromatin was counterstained by Hoechst (blue). Dotted lines show cell boundaries. Scale bar 5 µm. aMTs: astral microtubules. Bottom row: zoomed-in details (Scale bar: 2 µm). g) Differentiating cells increase their number of astral microtubules (as determined in f). Each data point represents a single cell (ESCs n = 115 cells, differentiation n = 122 cells from N = 3 experiments), the large circles denote the medians in each cell volume bin, the error bars show the interquartile ranges.

### Spindles switch to pronounced astral architectures in early-differentiated cells

Cells can change spindle size by changing the balance between microtubule nucleation, growth, and turnover, which is known to alter microtubule polymer mass and spindle volume ^9,53,54^. To test whether spindles in neurally-differentiating cells change their size because of altered microtubule dynamics, we measured microtubule growth velocities using the fluorescent plus end-tracking protein EB1::tdTomato (Fig. 2a). We acquired time-lapse recordings of undifferentiated ESCs and early-differentiated cells and tracked growing microtubule plus ends (Suppl. Movie 3). We found growth velocity to be independent of spindle size and mitotic cell volume (Suppl Fig. 3a) and identical between the differentiation states (v_p_ESC_ = 0.26 ± 0.034 µm/s, v_p_DIF_ = 0.26 ± 0.031 µm/s, Fig. 2b). Consistently, the cellular levels of the microtubule polymerase CKAP5 remained comparable (Suppl. Fig. 3b). Thus, despite the difference in spindle size, the microtubule growth dynamics remained unchanged.

**Figure 3:**
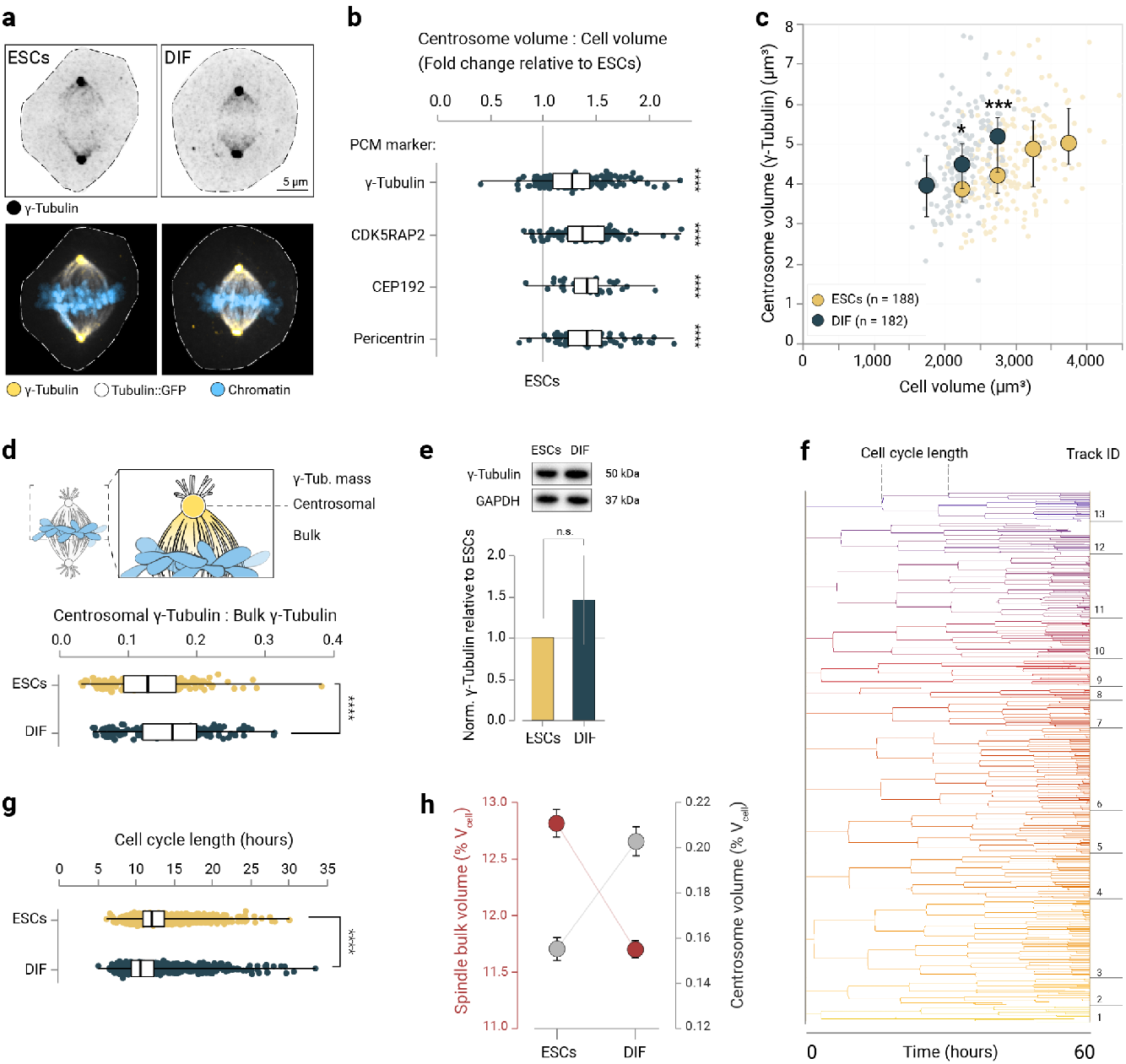
Centrosomes superscale upon differentiation. a) Confocal micrographs (maximum-projected) of fixed ESCs or early-differentiated cells at metaphase. Top: Immunostained γ-tubulin signal (inverted grayscale), bottom: immunostained γ-Tubulin (yellow), Tubulin::GFP (grey) and chromatin counterstained by Hoechst (blue). Dotted lines indicate the cell boundaries. Scale bar: 5 µm. b) The mitotic PCM is enlarged upon differentiation. Fold change in centrosome occupancy (Centrosome volume : cell volume) in early-differentiated ESCs relative to undifferentiated ESCs, comparing 4 centrosomal markers. Data points represent individual cells, the boxes show the interquartile ranges, the vertical lines show the medians and the whiskers show the extrema. Significances against the undifferentiated ESCs were tested with Welch’s t-tests, ****: p < 0.0001. c) Centrosome volume scales with cell volume. Centrosome volume based on γ-Tubulin. Each data point represents a single cell (ESCs n = 188 cells and differentiation n = 182 cells from N = 6 experiments), the large circles denote the medians in each cell volume bin (bin size = 500 µm^3^, the error bars show the interquartile ranges. Statistics per cell volume bin by Welch’s t-test, *: p < 0.05, ***: p < 0.001. d) γ-Tubulin mass is redistributed in early-differentiated cells. Top: γ-Tubulin localisation in centrosomes or within the spindle bulk. Bottom: centrosomal to bulk γ-Tubulin mass ratio in immunostained cells (see a). Data points represent individual cells (sample sizes as in c), the boxes show the interquartile ranges, the vertical lines show the medians and the whiskers show the extrema. Welch’s t-tests, ****: p < 0.0001. f) γ-Tubulin expression is comparable between the differentiation states. Top: representative western blots against γ-Tubulin in undifferentiated ESCs or early-differentiated cells. GAPDH as loading control. Bottom: γ-Tubulin (normalised to GAPDH), relative fold change. Bars show the mean (N = 3), error bars show the standard deviation. Welch’s t-tests, n.s: not significant, p > 0.05 g) Cell cycle length was determined by tracking of individual cell families. Visual representation of tracking data of an exemplary adaptive feedback recording (see Figure 1) of undifferentiated ESCs. h) Cell cycle length changes upon differentiation ^87,88^. Intermitotic times from time-lapse recordings (N = 9, see Figure 1) of undifferentiated ESCs (yellow, n = 2,614) or differentiating ESCs (blue, n = 2,230). Data points represent individual cells. The boxes show the interquartile ranges, the black lines inside the boxes denote the medians, whiskers show the extrema. Welch’s t-test, ****: p < 0.0001. i) As spindles subscale, centrosomes superscale upon differentiation. Comparing spindle bulk scaling to cell volume (data and sample sizes as in Fig. 1h) and centrosome scaling to cell volume (as determined by γ-Tubulin, data and sample sizes as in Fig 3c) between ESCs and early-differentiated ESCs). Circles show the means, error bars show the 95% confidence intervals.

However, we found a considerable switch in spindle architecture: at 48 hours of differentiation, microtubules increasingly grew from spindle poles but less in the spindle bulk (Fig. 2c, d and Suppl. Fig 3a). This led to a shift in the ratio of astral to spindle bulk microtubules (Fig. 2e) at a constant number of microtubules per cell volume (Suppl. Fig. 3c). To complement the live data, we additionally visualised microtubule populations via 3D immunofluorescence (Fig. 2f), which confirmed that the number of astral microtubules was significantly higher in early-differentiated cells when compared to undifferentiated ESCs of equal cell volumes (Fig. 2g). Intriguingly, in ESCs the number of astral microtubules seemed to saturate at a critical cell volume above ∼3000 µm^3^ (Fig. 2g). Together, these data indicated that the observed cell state-specific spindle scaling resulted from a redistribution of microtubules towards the asters away from the spindle bulk.

The switch in spindle architecture is analogous to observations in the developing mouse neocortex, where early neurogenic spindles displayed pronounced astral microtubules ^22^. In this system, TPX2, a MAP that stabilises microtubules and stimulates Augmin-mediated microtubule nucleation ^55–57^, has been identified as a main contributor of spindle morphology switches. In our system, however, neither cellular TPX2 nor Augmin levels, nor their localisation on spindles changed between undifferentiated ESCs and early-differentiated cells (Suppl Fig. 4). Thus, we concluded that the switch in spindle architecture was not a consequence of diminished TPX2- or Augmin-loading upon differentiation. Alternatively, microtubule number could be modulated by microtubule severing ^58–60^. Indeed, we found differences in Katanin localisation between stem cells and differentiating cells (Suppl. Figs. 5a-d). However, while Katanin knock-down did produce aberrant spindle phenotypes with buckled microtubules within the bulk (Suppl. Figs. 5e-h) ^61^ it had no effect on spindle volume (Suppl. Figs. 5i, j), ruling out microtubule severing as the main factor of spindle subscaling in our system. Taken together, these data suggested that differentiating cells changed their spindle architecture and size by redistributing microtubule growth to the spindle poles. This resulted in smaller spindles relative to cell size when comparing differentiating and undifferentiated cells.

**Figure 4:**
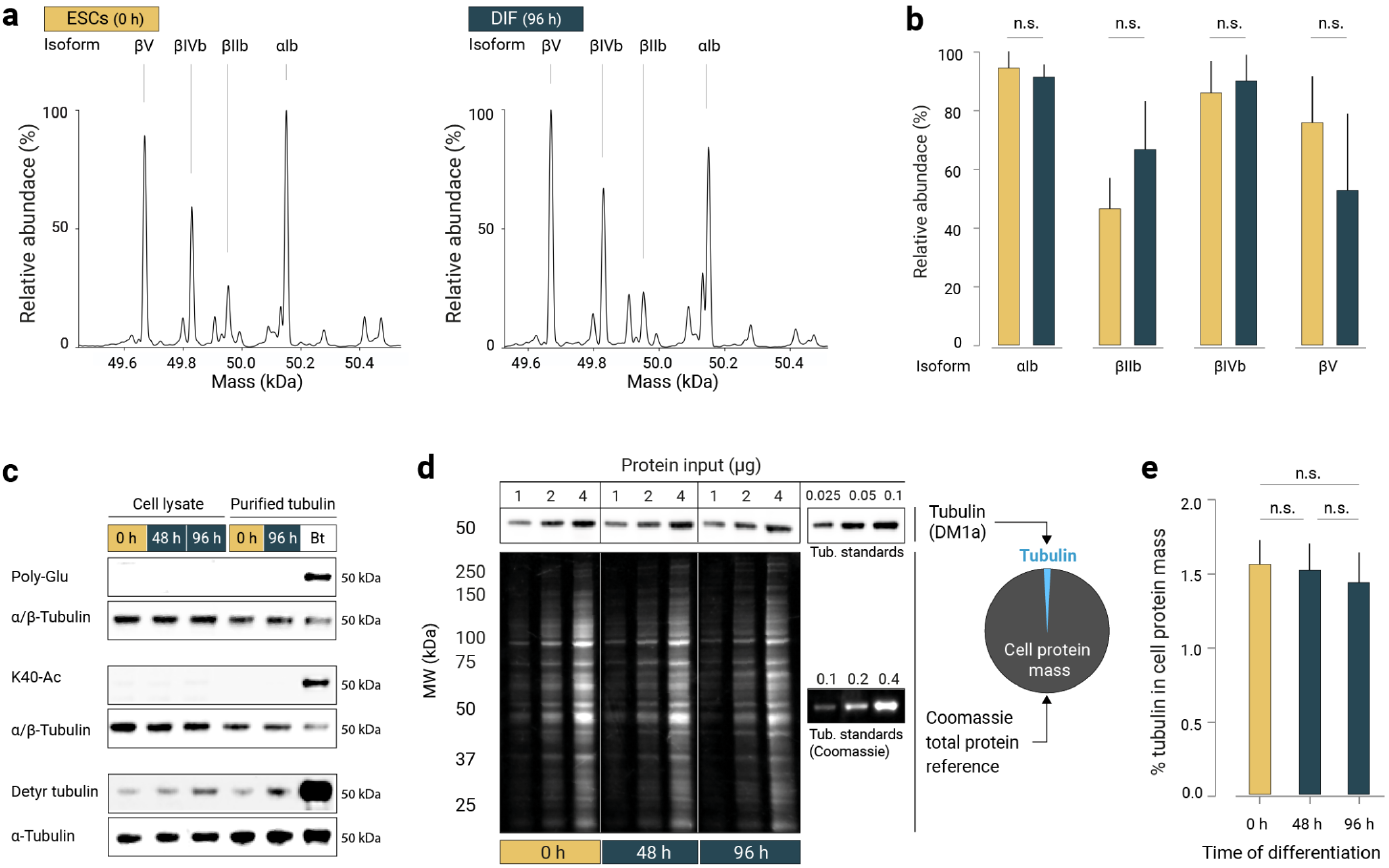
Tubulin biochemistry remains unchanged between the differentiation states. a) Deconvoluted mass spectra of tubulins purified from undifferentiated ESCs (left) or early-differentiated cells (right). Measured average masses of the most abundant signals and corresponding tubulin isoforms are labelled. Measured masses are in agreement with the theoretical masses of βV (MW = 49670.3 Da; Uniprot: P99024), βIVb (MW = 49830.5 Da, Uniprot: P68372), βIIb (MW = 49952.6 Da; Uniprot: Q9CWF2), αIb (MW = 50151.1 Da, Uniprot: P05213). b) The mean relative abundances of the 4 most dominant isoforms in N = 3 affinity-purified tubulin batches (ESCs vs. 96h-differentiated cells) measured by intact protein mass spectrometry. Error bars show standard error of the mean. Welch’s t-test, n.s.: not significant, p > 0.05. c) Tubulin post-translational modifications (PTMs) are comparable between the differentiation states. Western blots against various tubulin PTMs, using whole cell lysates and purified tubulin samples. Affinity-purified Bos taurus (Bt) brain tubulin was loaded as positive control. Poly-Glu: Poly-glutamylation. K40-Ac: Tubulin lysine 40 acetylation. Detyr: Detyrosinated tubulin. d) Top: Representative tubulin western blot of three differentiation timepoints (0 h, 48 h and 96 h) and whole-cell lysates (1 µg, 2 µg, 4 µg, N = 3). For downstream calibration, defined masses of purified tubulin were blotted onto the same membrane (N = 3 blots per batch of cell lysates). Bottom: Fluorescent detection of Coomassie-stained whole protein content on a replica gel. Again for downstream calibration, defined masses of purified tubulin were loaded. e) Tubulin consistently represents 1.5% of the cellular protein mass. Relative tubulin content in total cellular protein mass in whole-cell lysates of undifferentiated ECSs (0 h) versus 48 h or 96 h early-differentiated cells. Bars represent the mean, error bars show the standard error of the mean. Significances were tested with one-way analysis of variance (ANOVA), n.s.: Not significant.

**Figure 5:**
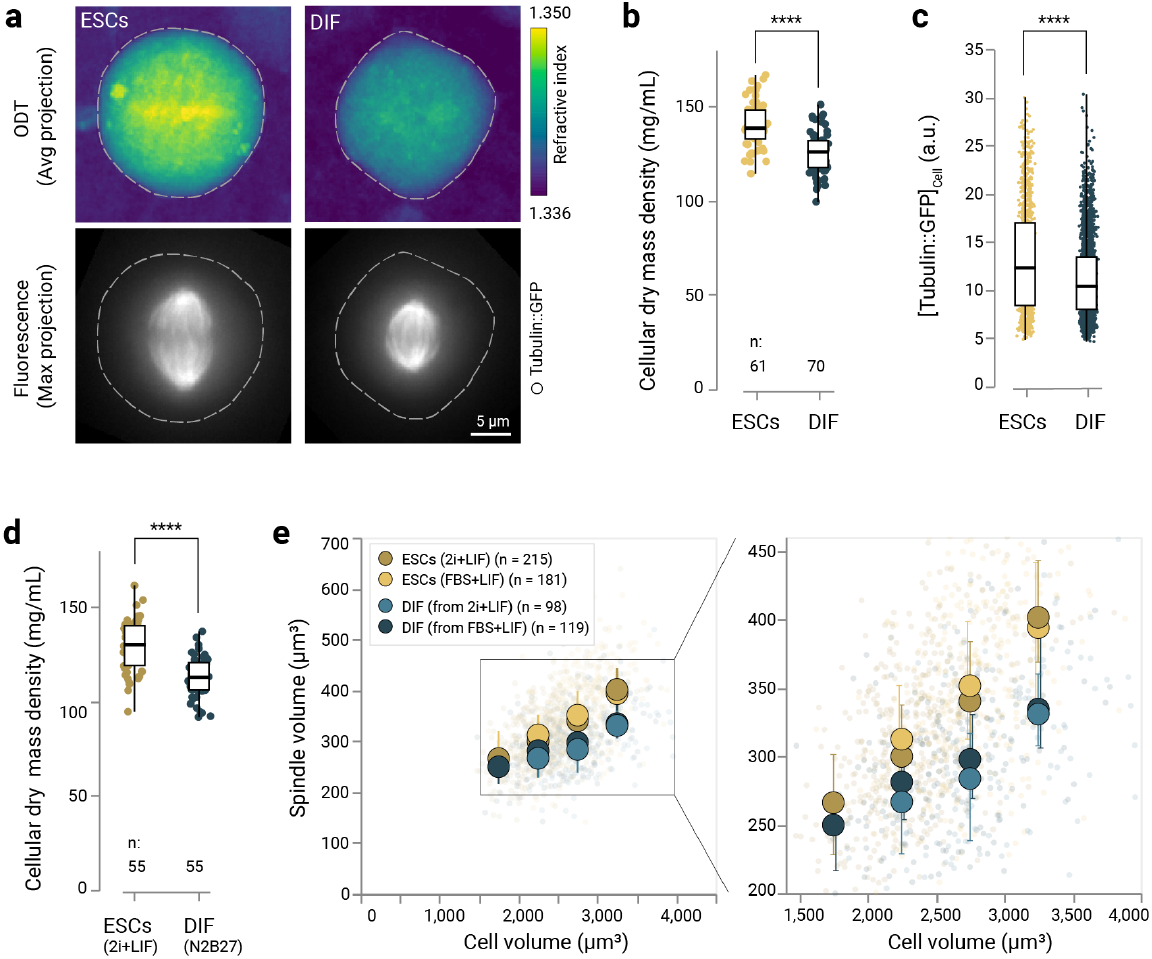
The cytoplasm is diluted in early-differentiating cells. a) Top: average projections of 3D refractive index maps derived from optical diffraction tomography (ODT) imaging of mitotic undifferentiated ESCs or early-differentiated cells. Colour coded according to the refractive index. Bottom: Maximum-projected epi-fluorescent micrographs showing the tubulin::GFP signal. Dotted lines show cell boundaries. Scale bars: 5 µm. b) Cellular mass density decreases upon differentiation. 3D cellular mass densities of mitotic undifferentiated ESCs versus early-differentiated cells. Data points show individual cells (ESCs n = 61 and differentiation n = 70 from N = 3 experiments). The boxes show the interquartile ranges, the black lines inside the boxes denote the medians, whiskers show the extrema. Welch’s t-test, ****: p < 0.0001. c) The average tubulin::GFP signal (3D total cell) decreases during differentiation. Data from long-term automated imaging (see Figure 1). Boxes show the interquartile ranges, horizontal lines show the medians, whiskers show the extrema. Data points show individual cells (ESCs n = 1,088 and differentiation n = 2,921 from N = 9 experiments). Significance tested by Welch’s t-test, ****: p < 0.0001. d) As in (b) but showing differentiated cells from 2i+LIF ESCs cultures (ESCs n = 55 and differentiation n = 55 from N = 2 experiments). Significance tested by Welch’s t-test, ****: p < 0.0001. e) Spindle volume subscaling is differentiation-intrinsic. Left: Comparing spindle bulk volume scaling to cell volume in early-differentiating cells originating from 2i+LIF ESCs cultures or from FBS+LIF ESCs cultures, as determined by spinning-disk confocal live cell imaging. Data points show individual cells (ESCs (2i+LIF) n = 215, ESCs (FBS+LIF) n = 181, DIF (from 2i+LIF) n = 98, DIF (from FBS+LIF) n = 215, pooled from N = 5experiments). Large circles represent the medians in 500 µm^3^ cell volume bins. Error bars show the interquartile ranges. Right: Zoomed-in detail.

### Centrosomes superscale in early-differentiated cells

The apparent increase in nucleation capacity at the spindle poles could simply be a result of increased centrosome size ^62^. Centrosomes are the major microtubule nucleating centers in mammalian cells and important for mitotic spindle assembly, asymmetric cell division, and cell polarity ^63,64^. Therefore, we directly visualised and measured centrosomes by staining γ-Tubulin (Fig. 3a) and components of the pericentriolar material (PCM), i.e. CDK5RAP2, CEP192, and Pericentrin (Suppl. Fig. 6a-c). We found that centrosomes superscaled upon differentiation, i.e. the relative volume of all centrosomal proteins increased by approximately 1.4-fold in early-differentiated cells when compared to the undifferentiated stem cells (Fig. 3b). Centrosome volume scaled with cell volume for both cell states (Fig. 3c), as has been observed in early *C. elegans* and Zebrafish embryogenesis ^4,65,66^. However, centrosomes of undifferentiated ESCs became independent of cell size and approached an upper limit in cells with volumes above ∼3000 µm^3^ (as measured by γ-Tubulin, Fig. 3c, and by CDK5RAP2, CEP192, and Pericentrin see Suppl. Fig. 6d-f). Indeed, in early-differentiated cells, more γ-Tubulin was recruited to the centrosomes when compared to stem cells (Fig. 3d), while overall γ-Tubulin levels did not change (Fig. 3e). The relative increase in centrosome size paralleled the increase in nucleation capacity and astral microtubule number in early-differentiated cells (see Fig. 2g). This suggested a direct relation between centrosome volume and nucleation capacity, as has been reported previously ^62^.

**Figure 6:**
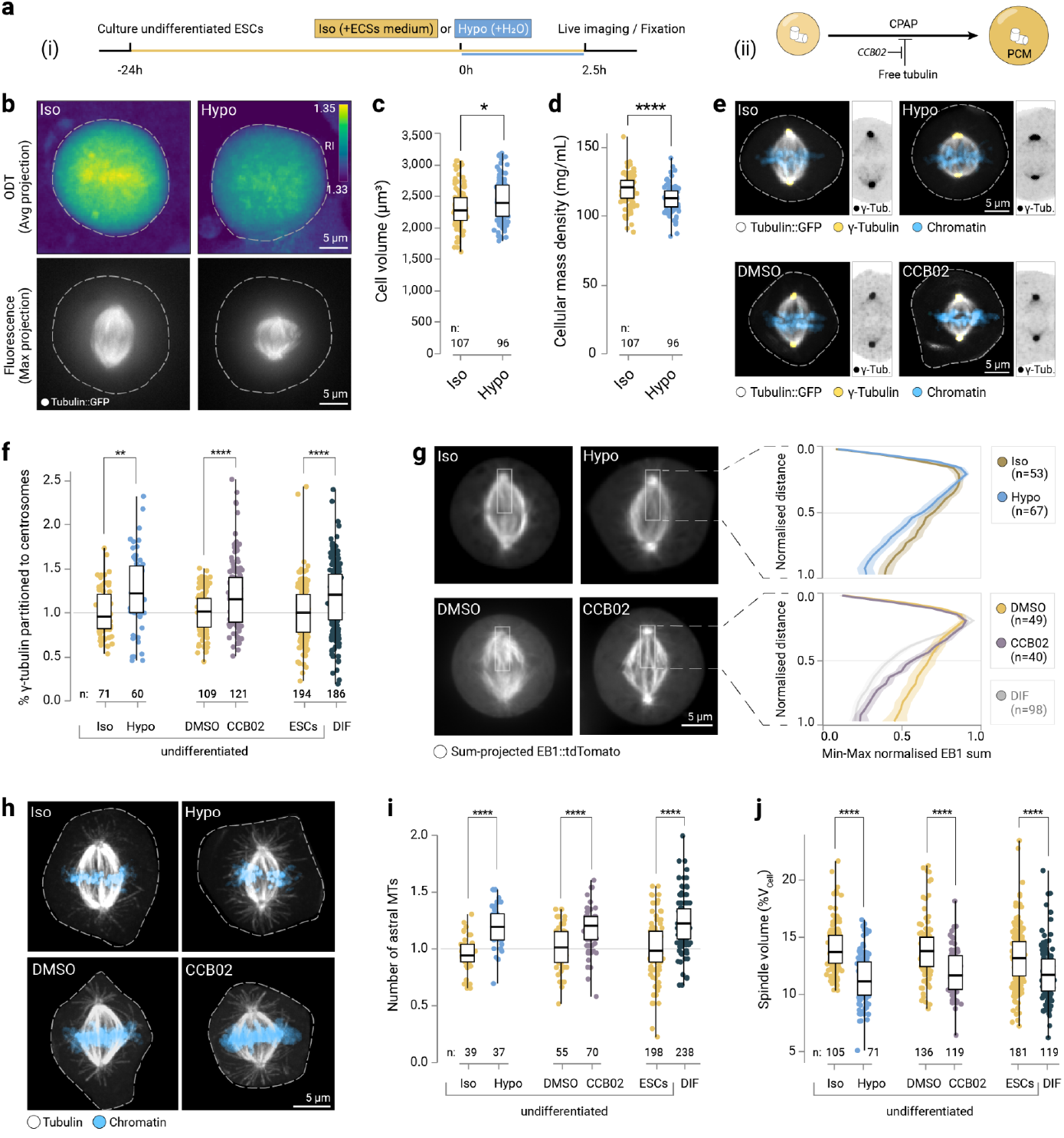
Cytoplasmic dilution shifts spindle architecture by liberating CPAP and increasing centrosomal nucleation capacity. a) Strategies for osmotic perturbation of undifferentiated stem cells (i) and small molecule inhibitor treatment liberating CPAP from its inhibitory binding to tubulin in undifferentiated stem cells (ii). b) Top: average projections of 3D refractive index maps derived from ODT imaging of mitotic ESCs, with 2 h recovery after adding isotonic medium (Iso, 337 mOsmol/kg) or ultra pure water (Hypo, 250 mOsmol/kg). Colour-coded according to the refractive index. Bottom: Maximum-projected epi-fluorescent micrographs showing the tubulin::GFP signal. Dotted lines show cell boundaries. Scale bars: 5 µm. c) Cells swell upon hypoosmotic challenge. ODT-derived cell volumes after hypoosmotic treatment of ESCs. Boxes show the interquartile ranges, horizontal lines show the medians, whiskers show the extrema. Data points show individual cells (n isosmotic: 107, n hypoosmotic: 96 from N = 4 experiments). Significance tested by Welch’s t-test, *: p < 0.05. d) As (c) but showing mitotic cell mass density. ****: p < 0.0001. e) Maximum-projected confocal micrographs showing immunostained γ-Tubulin signals (yellow, or inverted grey (cropped images), respectively) in fixed ESCs, Tubulin::GFP in grey, Hoechst-stained chromatin in blue. Top: Cells after hypoosmotic challenge (“Hypo”) versus control (“Iso”), bottom: cells after CCB02 treatment versus control (“DMSO”). Scale bars: 5 µm. f) The fraction of total γ-Tubulin signals residing at the centrosomes, comparing iso- (n = 71) versus hypoosmotically-treated ESCs (n = 60) (N = 2 experiments, normalised to the isosmotic control), and comparing DMSO- (n = 109) versus CCB02-treated (n = 121) ESCs (N = 3 experiments, normalised to the DMSO control), and comparing undifferentiated ESCs (n = 194) with early-differentiated ESCs (n = 186) (N = 6 experiments (see Figure 3b), normalised to undifferentiated ESCs). The boxes show the interquartile ranges, the black lines inside the boxes denote the medians, whiskers show the extrema. Significance tested by Welch’s t-test, **: p < 0.01, ****: p < 0.0001. g) Left: Sum-projected movies (20 s) of mitotic ESCs labelled with EB1::tdTomato, after iso- or hypoosmotic treatment (top), or after treatment with CCB02 (10 µM) or DMSO-treated as control (bottom). Right: Min-max normalised half-spindle intensity profile plots of the sum-projected EB1::tdTomato movies of iso-(n = 53) or hypoosmotically treated (n = 67) ESCs (N = 2 experiments), and of ESCs after CCB02 treatment (n = 40) (versus DMSO control, n = 49) (N = 2 experiments). For reference, the profile of early-differentiated cells (data as in Fig. 2d) is shown. The lines show the means and the shaded areas show the 95% confidence intervals. Dotted lines mark the norm. EB1 intensity at the midpoint distance. Welch’s t-test, *: p < 0.05, **: p < 0.0. h) Confocal micrographs (max-projected) of metaphase control (“Iso”) ESCs or after hypoosmotic treatment (“Hypo”) (top row) or CCB02-treated or DMSO-treated ESCs (bottom row), fixed and stained with anti-tubulin antibodies (grey). Chromatin is stained with Hoechst (blue). The dotted lines show cell boundaries. Scale bar: 5 µm. i) As in (g) but showing the number of astral microtubules, comparing iso-(n = 39) versus hypoosmotically treated ESCs (n = 37) (N = 2 experiments, normalised to the isosmotic control), and comparing DMSO- (n = 55) versus CCB02-treated (n = 70) ESCs (N = 2 experiments, normalised to the DMSO control), and comparing ESCs (n = 198) with 48 h-differentiated ESCs (n = 238) (N = 3 experiments (same data as in Figure 2g), normalised to ESCs). j) As in (g) but showing the percentage of cell volume occupied by the spindle, comparing iso- (n = 105) versus hypoosmotically treated ESCs (n = 71) (N = 2 experiments), and comparing DMSO-(n = 136) versus CCB02-treated (n = 119) ESCs (N = 3 experiments), and comparing ESCs (n = 181) with 48 h-differentiated ESCs (n = 119) (N = 5 experiments (same data as in Figure 5e)).

Commonly, to prepare for mitosis, microtubule remodelling starts with a dramatic increase in microtubule nucleation at the centrosomes driven by the recruitment and local activation of γ-TuRC ^62,67^. To ensure that the enlarged centrosomes observed in early-differentiated cells were not simply persisting centrosomes from larger stem cells, we used our long-term imaging strategy to track individual cells across several generations (Fig. 3f). This confirmed that cells classified as early-differentiated cells had a cell cycle duration of ∼11 hours and divided 3 to 4 times (Fig. 3g), implying they dynamically remodelled their PCM. Taken together, these data suggested that the relative increase in centrosome size shifts the microtubule nucleation capacities towards the spindle poles at the expense of the spindle bulk (Fig. 3h) leading to smaller spindles with larger asters in early-differentiated cells.

### Tubulin biochemistry remains unchanged between the differentiation states

Another way of changing spindle architecture in differentiating cells could be through fundamental changes in tubulin biochemistry. Therefore, we measured tubulin isoform composition, tubulin post-translational modifications (PTMs), and tubulin levels. To determine tubulin isoforms and PTMs, we purified tubulin from either ESCs or early-differentiated cells ^68^. Intact protein mass spectrometry revealed no significant changes in isoform composition between day 0 and 4 of differentiation (Fig. 4a, b). Significant βIII-Tubulin expression (as a marker of neurons) started only with terminal differentiation later than day 4 (Fig. 1a) as reported previously ^47^. Similarly, there were no differences

between the PTM patterns in ESCs and early-differentiated cells (Fig. 4c). Together, this showed that neither tubulin isoform composition nor tubulin PTMs change during early differentiation. Next, we measured cellular tubulin levels by quantitative immunoblots (Fig. 4d, see Methods). For each time point during differentiation, tubulin constituted approximately 1.5% of total cellular protein mass (Fig. 4e). From these data we conclude that changes in spindle morphology are not driven by tubulin biochemistry, and thus other spindle scaling mechanisms must be operating.

### The cytoplasm is diluted in early-differentiating cells

Differentiation-induced changes in cell function have been correlated with reductions in cellular mass density in a variety of cellular systems ^34–39^. To test whether in our system cells change their dry mass density (defined as dry mass/volume, see Methods) ^29^ during differentiation, we measured subcellular distributions of refractive indices using correlative fluorescence and optical diffraction tomography ^69^. Remarkably, we found that cellular dry mass density in early-differentiated cells was reduced by 10% relative to the undifferentiated ESCs (ρ_ESC_ = 140 ± 12 mg/mL vs ρ_DIF_ = 125 ± 11 mg/mL, Fig. 5a, b). This differentiation-associated dilution of the cytoplasm was observed across all cell volumes (Supplementary Fig. 7). Consistently, we measured a reduction in total cellular (spindle and cytoplasmic) tubulin::GFP fluorescence using live-cell imaging (Fig. 5c). This indicated a drop of total tubulin concentration concomitant with cytoplasmic dilution. The reduction in total tubulin did not lead to a reduction in microtubule density within the spindle bulk (see Suppl. Figs 2f, 3a). Importantly, both the observed decrease in cellular mass density as well as changes in spindle scaling were independent of culturing and differentiation conditions (Fig. 5d, e). Taken together, these data imply that changes in cellular mass density could have consequences for spindle architecture and scaling during differentiation.

### Cytoplasmic dilution shifts spindle architecture by liberating CPAP and increasing centrosomal nucleation capacity

The above data suggested that tuning cellular mass density could contribute to changes in spindle volume. If cytoplasmic dilution altered spindle morphology universally, we would expect a differentiation-independent drop in cellular mass density to affect spindle size in undifferentiated ESCs. To test this, we diluted the cytoplasm of stem cells by lowering the osmolality of the culturing medium (Fig. 6a, b). As expected, a decrease in osmolality led to an increase in cell volume and a reduction in cellular mass density (ρ_ISO_ = 118 ± 10 mg/mL vs ρ_HYPO_ = 112 ± 9 mg/mL, Fig. 6c, d). This drop in cellular mass density led to an increase in γ-Tubulin at the centrosomes (Fig. 6e, upper panel and f) and consequently an increase in centrosomal EB1 concentration (Fig. 6g) and higher numbers of astral microtubules (Fig. 6h, upper panel and i). Taken together, these data confirmed that a reduction in cellular mass density is sufficient to increase the centrosome’s nucleation capacity at the expense of the spindle bulk (Fig. 6j). These data imply that the spindle assembly mechanism is intertwined with the physical properties of the cytoplasm. But how is this implemented at the molecular level?

So far, there is no established link between cytoplasmic dry mass density and centrosome function. However, it has been reported that centrosomes can increase their nucleation capacity when free cytoplasmic tubulin decreases. This is because of the inhibitory interaction of soluble tubulin with the centrosomal protein CPAP ^70^. We speculated that the observed decrease in free tubulin liberated CPAP, which in turn increased PCM recruitment and centrosomal microtubule nucleation. To mimic a decrease in free tubulin in stem cells, we liberated CPAP from tubulin using a small molecule inhibitor (CCB02 ^71^, Fig 6a). This should result in an increase in astral microtubule number and a decrease in spindle bulk without changing cell size. Indeed, upon inhibitor treatment, spindles in undifferentiated ESCs partitioned more γ-Tubulin to the centrosomes (Fig. 6e, bottom panel, and f), which showed an increase in nucleation capacity as evidenced by a higher centrosomal EB1 concentration (Fig. 6g) and astral microtubule number (Fig. 6h, bottom panel, and i), leading to a reduction in spindle volume in equally sized cells (Fig. 6j). Thus, we propose that CPAP is a molecular determinant of centrosomal nucleation capacity that is responsive to cytoplasmic dilution. Together, our physical perturbation via cytoplasmic dilution and biochemical perturbation using a small molecule inhibitor allowed us to phenocopy the spindle architecture characteristic of early-differentiated cell states in undifferentiated stem cells.

Importantly, our data are consistent with previous quantitative models of spindle scaling ^9,18,53^. In the Supplementary Note, we provide a theoretical model that allows us to link cell volume, spindle volume and cytoplasmic density to estimate the number of astral microtubules. In our model, through the inhibition of a centrosomal regulator by free tubulin, the astral microtubule number has a Michaelis-Menten dependence on cell volume and is proportional to the concentration of free CPAP. This model is able to reproduce our experimental observations in stem cells and differentiating cells (Fig. 7a). Interestingly, this shows that spindle architecture can change significantly while total microtubule number does not. To ensure spindle size regulation during differentiation, this separation allows fine-tuning of spindle architecture without altering gross microtubule dynamics.

**Figure 7:**
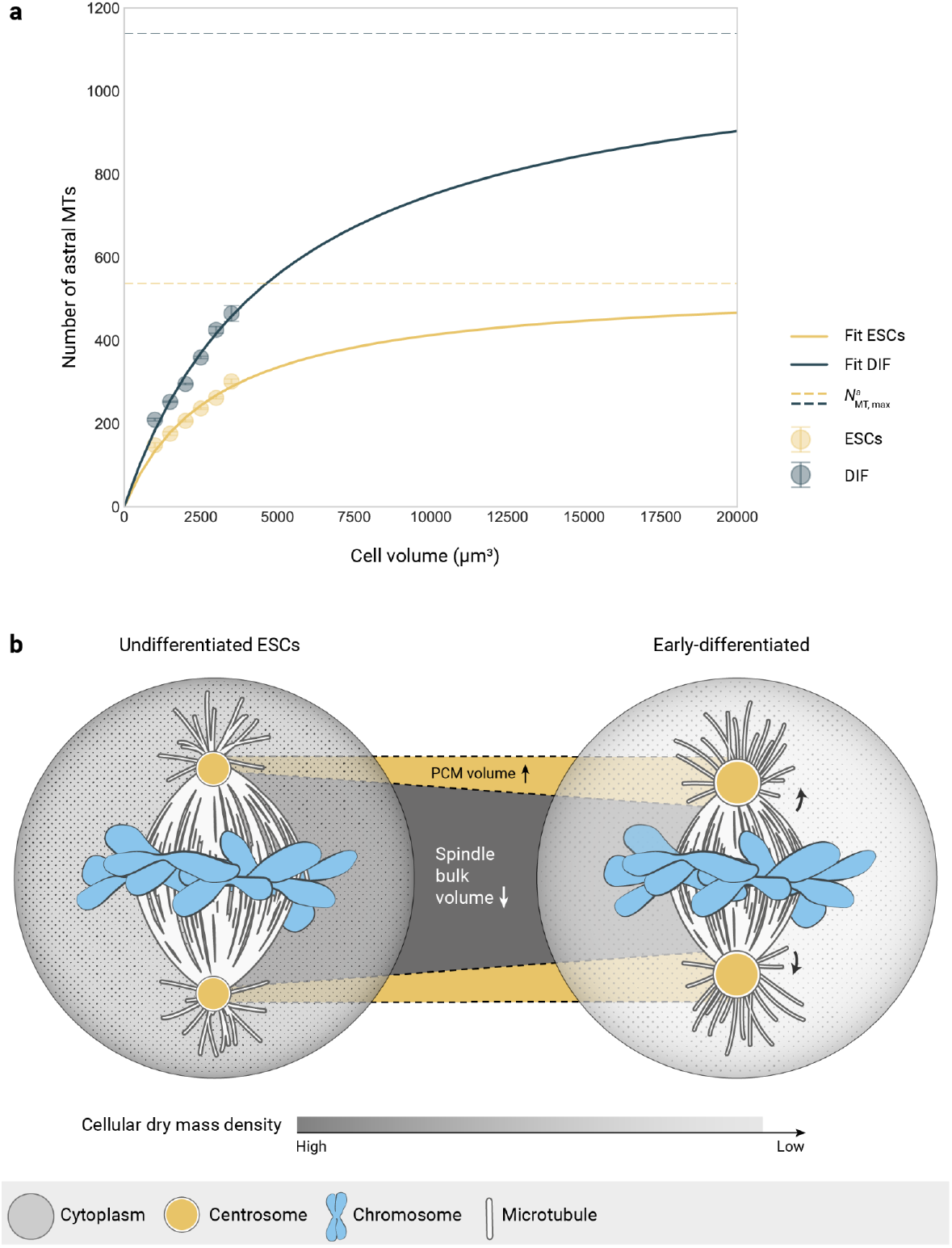
Cytoplasmic dilution-driven changes in mitotic architecture in early-differentiated cells. a) Fit of the theory to experimental data. For both cell states (undifferentiated: yellow; differentiated: blue), the number of astral microtubules is estimated from the spindle volume data (see theory) and grouped by cell volume in bins of 500 µm^3^. The large circles depict the averages for each cell volume bin and are fitted to equation (6), the error bars show the standard deviation. Solid curves represent the fit curves. Dashed lines depict the estimated saturation value for astral microtubule number. b) Upon differentiating, a reduction in cellular mass density leads to the enlargement of the pericentriolar material (PCM), increasing centrosomal nucleation capacity, and the redistribution of microtubule mass away from the spindle bulk towards the asters.

Thus, by combining our experimental data and theory, we propose that the fundamental physical property of the cytoplasm - its dry mass density - provides a mechanism for regulating spindle architecture and size during differentiation.

## DISCUSSION

The remarkable capacity of the spindle to scale with cell size has been observed across eukaryotes with spindle lengths ranging from 2 µm (yeast) to >50 µm (frog oocytes). Spindles showcase a large range in size not only between species but also within a single organism ^72^. Yet, how spindles adapt to changes in cell size, shape, and cytoplasmic properties as cells undergo differentiation was unclear. Here, we systematically quantify changes in spindle morphology and size during differentiation. Our data are consistent with a novel spindle scaling mechanism in which a reduction in cellular mass density leads to a relative increase in centrosome size in early-differentiated cells. As a consequence, microtubule nucleation shifts towards the spindle poles in these cells, promoting the generation of astral microtubules at the expense of the spindle bulk (Figure 7b). This mechanism explains how a fundamental physical property of the cell - dry mass density - can lead to changes in the spindle’s nucleation profile and consequently spindle architecture.

### Spindle scaling mechanisms

Based on the current knowledge of spindle assembly, a number of biophysical mechanisms has been established that explain the regulation of spindle size and the coupling to cell size to achieve spindle scaling. In the limiting component (volume-sensing) model, the finite availability of critical components has been implicated in the regulation of organelle size. In the context of spindle assembly, limiting components can be structural ^18,19^ or regulatory ^8,13^. A special case of component limitation is the cell surface-sensing mechanism, where the active cytoplasmic concentration of a spindle assembly factor is regulated through the sequestration of importin-α to the membrane ^6,9,20^. Both the volume-sensing and surface-sensing scaling models describe physical mechanisms where the coupling of spindle size to cell size is inherently self-correcting. Here, we propose that another physical property of the cell is essential to spindle size regulation. We propose that changes in cellular dry mass density explain the cell state-specific scaling phenotypes in a differentiating system (Figure 7). Indeed, a relationship between decreasing mass density and progressing differentiation has been proposed in many systems, including chondrocytes, keratinocytes, myeloid precursors, and neurons ^34–39^. Still, it is not immediately evident why and how neurally-differentiating cells would change their mass density, in particular before terminal differentiation. One idea is that single-cell properties affect the overall mechanical properties of the developing tissue and this - in turn - influences differentiation ^73–75^. This, however, was not the scope of this study and will therefore be part of future investigations.

In contrast to early stages of development, which typically are transcriptionally silent ^76,77^, differentiation is not. Why would a physical mechanism for spindle scaling be in place? Cell-state transitions are often asynchronous within a tissue ^78–80^. Thus, programmed scaling mechanisms based only on developmental timing or cytoplasmic composition would not robustly couple spindle size to cell size, potentially leading to errors in spindle positioning and chromosome segregation. This might be particularly relevant in neural development, a process that is notoriously sensitive to mutations in spindle and centrosome genes ^42,43^.

We find that CPAP is a molecular determinant of centrosomal nucleation capacity that is responsive to cytoplasmic dilution. Mutations in the CPAP gene link centrosomal defects to primary autosomal recessive microcephaly, a disorder characterised by severely reduced brain size and cognitive disability ^81^. Consistently, mitotic spindle orientation defects have been described in cells depleted of CPAP ^82,83^. While a direct link between microcephaly and spindle orientation defects remains to be definitely established, our data provide an explanation how interfering with CPAP function can have consequences for spindle architecture. Moreover, our findings are consistent with studies in neural stem cells of the embryonic developing mouse neocortex. Here, metaphase spindles displayed differences in their architecture from early to late neurogenic stages. At early neurogenic stages (that is, comparable to the early-differentiated cells in our system), spindles displayed long astral microtubules and a reduced microtubule density near the chromosomes ^22^. Together, these observations suggest that centrosome function via CPAP and resulting switches in spindle architecture could be a potential source for major developmental defects.

### Coordination of microtubule nucleation pathways

Mitotic spindle assembly mainly relies on two pathways that generate microtubules: centrosome- and chromatin-driven microtubule nucleation. Both pathways depend on the recruitment of γ-Tubulin, either to the centrosome or to pre-assembled microtubules. The current understanding is that in very large cells, spindle scaling is mainly achieved via microtubule nucleation to create sufficient amounts of polymer mass, while in small cells

regulating microtubule dynamics can be sufficient to scale spindle size ^9^. Indeed, the dominance of nucleation pathways has been shown to shift in the course of mitosis ^84^ or even gradually over the course of development ^6,10^. This cooperation of pathways is thought to provide robustness and mitotic fidelity ^85^, but as we show here, can also result in the construction of spindles with differing architectures, providing the necessary plasticity to adapt spindles to evolving cellular environments.

## Supplementary note

**Supplementary Note Figure 1:**
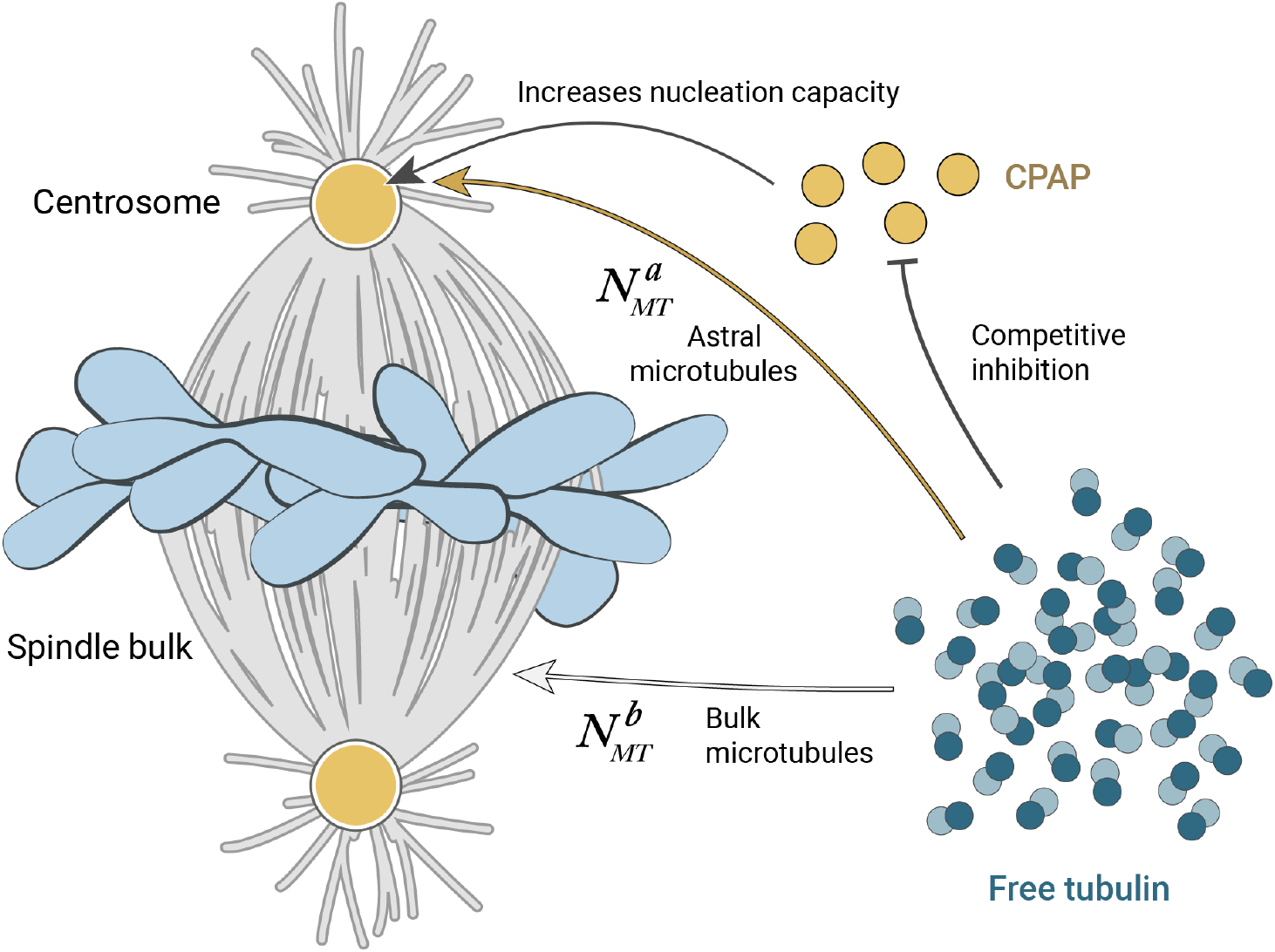
Schematic explaining the redistribution of mitotic microtubule populations via the inhibition of CPAP by free tubulin.

### Theoretical model

In this study, we have shown that differentiating cells change their spindle architecture and size by redistributing microtubule growth to the spindle poles at the expense of the spindle bulk (Suppl. Note Fig. 1). While the total microtubule number scales with cell volume (Suppl. Fig. 3c), the distribution into either astral or spindle bulk microtubules defines final spindle size. Thus, we propose a scaling of astral microtubule number with cell volume based on the mechanism suggested by Rieckhoff and colleagues (Rieckhoff et al. 2020): astral microtubule number has a Michaelis-Menten dependence on cell volume and is proportional to the number of active (centrosomal) nucleators in the cell:

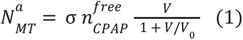

where 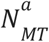 denotes the number of astral microtubules, *V* the cell volume and 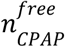 the concentration of free CPAP molecules (analogous to an active nucleator in Rieckhoff et al. 2020). The model parameters are σ and *V*_0_, which respectively denote the proportionality constant between the number of astral microtubules and the free CPAP concentration, and the reaction-diffusion volume of CPAP. The reaction-diffusion volume is defined as the volume in which the reaction is not limited by diffusion. The observed inhibition of CPAP by free tubulin (Fig. 6) is an important aspect of the system, which we incorporate into the theory in the following way: tubulin competitively inhibits CPAP through a simple binding reaction with a dissociation constant κ_*c*_. CPAP bound to tubulin is unable to increase centrosomal nucleation capacity, only free CPAP molecules can, their concentration being:

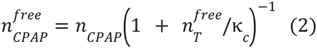

where *n*_*CPAP*_ denotes the total concentration of CPAP in the cell (free + bound to tubulin) and 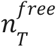 the soluble tubulin concentration in the cell. Tubulin is either soluble or polymerised, which can be expressed with the following equation:

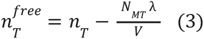

where *n*_*T*_ denotes the total tubulin concentration in the cell and 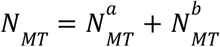 the total number of microtubules, which is the sum of the number of astral microtubules 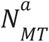 with the number of spindle bulk microtubules 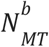. We combine the linear density of tubulin α, the average growth velocity *v*_*MT*_ and the average lifetime of a microtubule τ_*MT*_ in a single parameter λ (λ = α *v*_*MT*_ τ_*MT*_), which represents the number of tubulin molecules found in an average microtubule and which we assume to not significantly change between undifferentiated and differentiated cells based on our experimental evidence (Fig. 2).

To test whether our model can correctly predict a higher number of astral microtubules 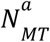 for differentiated cells, we connect both microtubule populations, i.e. the number of microtubules in the spindle bulk 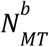 and astral microtubules 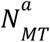, using our experimentally determined ratio *f*_*T*_ (Fig. 2e):

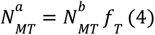

Thus, our model is fully described by equations (1)-(4) and these already qualitatively predict more astral microtubules and less bulk microtubules in differentiated cells when compared to undifferentiated cells with the same cell volume *V*.

### Model predictions

First, we establish which quantities of the model stay constant and which change when we compare undifferentiated and differentiated cells. As previously mentioned, our experimental results show that we can consider microtubule dynamics related quantities such as λ to be the same. The dissociation constant κ_*c*_ between tubulin and CPAP has been reported to be 3. 6 µM ^71^ and we assume it to have this value for both undifferentiated and differentiated cells. Lastly, we assume the proportionality constant σ between the number of astral microtubules 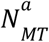 and the concentration of free CPAP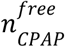to be the same for both systems. This assumption can be justified by the observation that in both cell states an increase in centrosome size correlates with nucleation capacity (Fig. 2g and 3c).

Quantities that we can expect to change across differentiation are those related to dry mass density, given that we observe an approx. 10% reduction of dry mass density in differentiated cells compared to undifferentiated cells (Fig. 5). In our model, the total concentration of CPAP *n* _*CPAP*_ and the total concentration of tubulin *n*_*T*_ are both reduced in differentiated cells by the same factor as the dry mass density (see Fig. 4, Fig. 5 and Suppl. Note Fig. 2). Finally, we can expect the reaction-diffusion volume of CPAP *V*_0_ to be larger for differentiated cells because of their lower dry mass density, which decreases crowding. With these considerations, we see from equation (3) that the free tubulin concentration 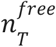 is lower in differentiated cells than undifferentiated cells of the same volume. Starting from equation (2), let us derive under which conditions the free CPAP concentration is 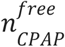 higher in differentiated cells. The expressions for both cell states are:

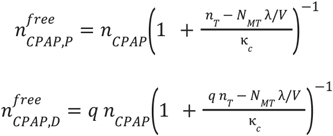

where the subindices P, D denote undifferentiated (pluripotent) and differentiated quantities, respectively. The factor 0 < *q* < 1 describes the decrease of the overall mass density in differentiated cells compared to undifferentiated cells. If we consider cells of the same volume, then the term *N*_*MT*_ λ/*V* is the same for both cell states. We can rearrange the expression for 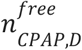 in the following way:

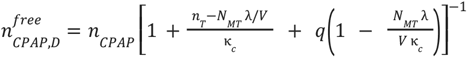

From the equation above we can see that the condition for 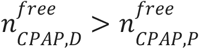 is:

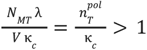

where we defined 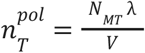 as the effective concentration of polymerised tubulin in the cell.

Based on our measurements of total cellular tubulin (Supplementary table S1), we estimate 50% of the total amount of tubulin to be in the polymer form 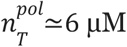, consistent with previous reports ^89^. As mentioned above, the dissociation constant κ _*c*_ = 3. 6 µM ^71^, so indeed 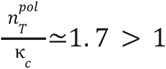. Therefore, by considering equation (1), we see that both the higher values of 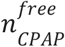 and *V*_0_ in differentiated cells will qualitatively result in a higher number of astral microtubules 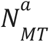.

**Supplementary Note Figure 2:**
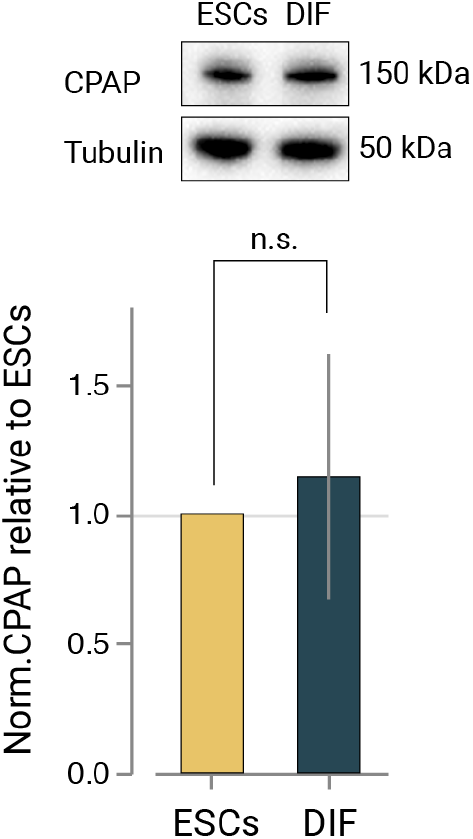
CPAP : total tubulin stoichiometry is unchanged between the differentiation states. Cellular levels of CPAP after 48 h of differentiation relative to the undifferentiated ESCs, probed by western blotting and normalised to tubulin (N = 3). Bars show the mean, errors show the standard deviation. Significance tested by Welch’s t-test, n.s.: p > 0.05.

### Model fitting

Now that we have shown that the model works qualitatively, we next want to connect it to our measured quantities. To estimate the number of microtubules in the spindle bulk, we link our experimentally well-defined spindle volumes *V*_*S*_ and tubulin concentration in the spindle bulk 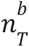 by

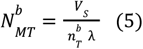

Once we determined 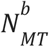, we can obtain 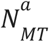 using equation (4). Note that we assume 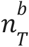 to be the same for undifferentiated and differentiated cells (based on Suppl. Fig. 2f, Suppl. Fig 3a, Suppl. Fig 4).

In order to fit our data with the model, we use equations (4) and (5) to estimate 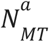 with our data for *V*_*S*_ and rewrite equation (1) as follows:

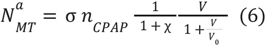

where 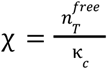 and *V*_0_ are treated as fit parameters. Next, we have to estimate the parameter σ, which collapses the influence of CPAP on centrosome size and nucleation capacity. To do so, we rearrange equation (1):

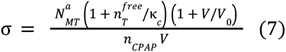

In order to estimate 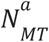 in the above equation, we can use experimentally measured spindle quantities:

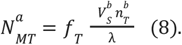

### Parameter values and estimates

All parameter values are given in Supplementary Note Table S1, more specifically the column that corresponds to cells binned by volume (V_bin_). For the purpose of estimating σ using equation (7), we need an estimate for *V*_0_, the reaction diffusion volume, and for 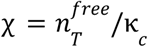. For these, we assign the following values: *V*_0_ = 3000 µm^3^, which is consistent with our data (Fig. 2g) and with both cell states being in the linear scaling regime ^5,9,72^. As described above, we estimated 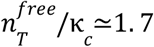 based on our measurements and a published κ_*c*_ ^71^. Plugging these values into equations (7) and (8) results in a value range σ = 0. 003∼0. 006. Therefore, we assume σ = 0. 0045 for both undifferentiated and differentiated cells in the following fits of the experimental data to the theory.

### Model results & discussion

To test our model, we take our estimates of 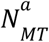 for both cell states, bin them by cell volume analogous to the experimental data, and fit the respective averages of 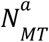 to equation (6).

Supplementary Note Figure 3a shows the binned averages for 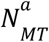, the respective fit curves, and the obtained fit parameters for both cell states. For differentiated cells, the fitted value of *V*_0_ is larger than for undifferentiated cells but lies in the same order of magnitude (10^3^ µm^3^).

This is consistent with a drop in cellular mass density and further implies that the number of astral microtubules will start to saturate with cell volumes > 10^3^ µm^3^. This is indeed what we observe for astral microtubules (Figure 2g) and centrosome volume (Figure 3c, Suppl. Fig. 6). For the fit parameter 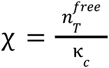 we can see that differentiated cells exhibit a lower value than undifferentiated cells. This again is consistent with a reduction in free tubulin concentration in differentiated cells. Importantly, the model gives a reasonable fit to the binned cell volumes as well as the respective point cloud of our experimental data (as presented in the inset plot of Supplementary Note Figure 3a). In Supplementary Note Figure 3b, we test how the fit parameters *V*_0_ and χ influence the number of astral microtubules as a function of cell size as defined by equation (6). Changing the parameter χ results in the same curve multiplied by a numerical prefactor. This is consistent with our biochemical perturbation experiments (see Figure 6), where we inhibit the CPAP tubulin interaction, which would decrease χ. The parameter *V*_0_ represents the cell volume at which the system reaches half of the saturation value, but it also affects the saturation value itself as we can see by considering equation (6) in the limit *V*/*V*_0_≫1:

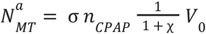

This is consistent with the number of astral microtubules as well as centrosome volumes reaching the saturation regime at different levels for both cell states (Fig. 2g and 3c).

A main point in our observation is that the total number of microtubules scales with cell volume and does not change between cell states (Suppl. Figs. 3a, c). To test whether our theory would correctly reproduce this observation, we lastly estimate the total number of microtubules. Indeed, we find the total number of microtubules to scale with cell volume and to not differ significantly between cell states (Suppl. Note Fig. 3c).

**Supplementary Note Figure 3:**
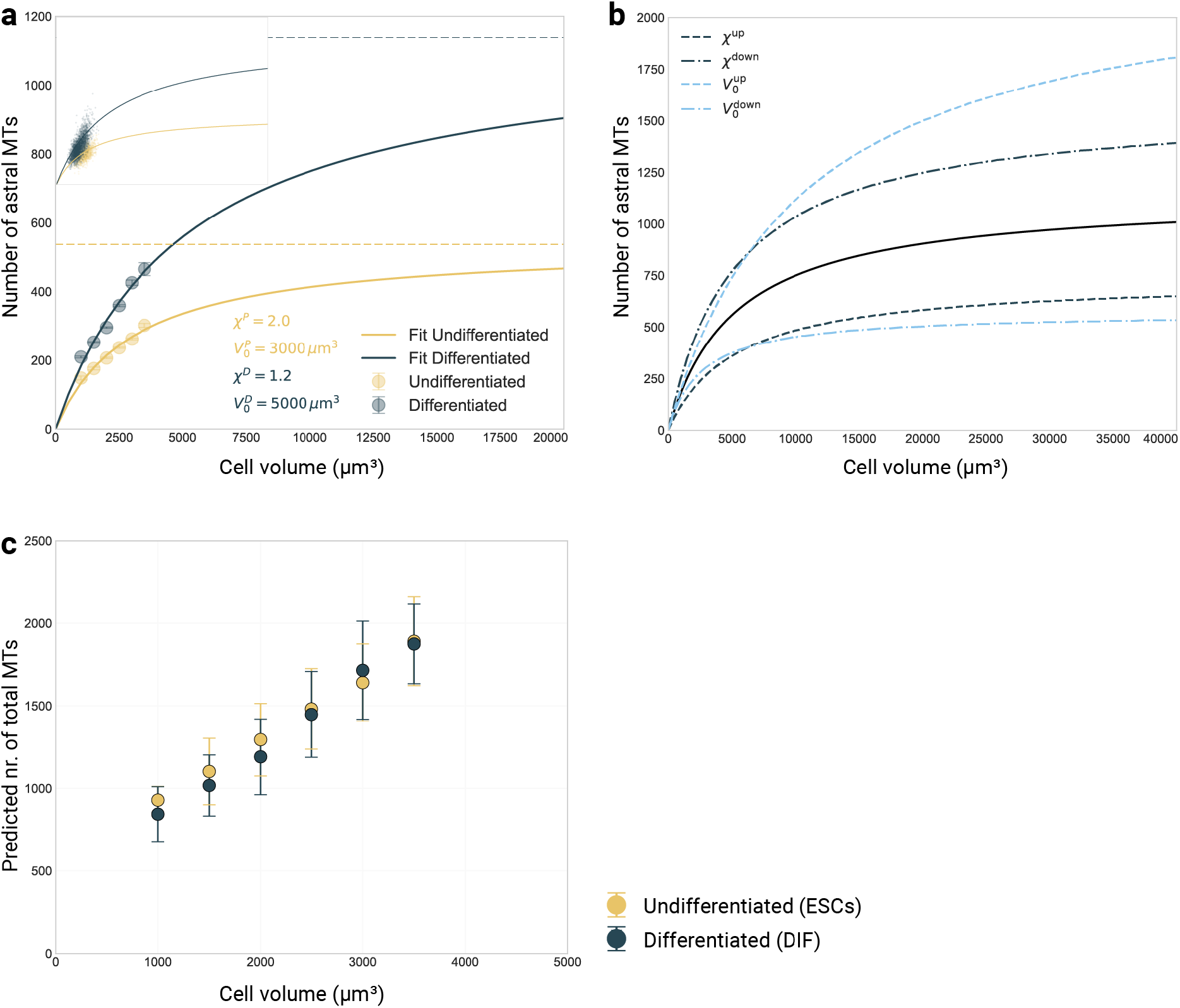
Results of the theoretical model. a) Fit of the theory to experimental data. For both cell states (undifferentiated: yellow; differentiated: blue), the number of astral microtubules is estimated from the spindle volume data and grouped by cell volume in bins of 500 µm^3^. The large circles depict the averages for each cell volume bin and are fitted to equation (6), the error bars show the standard deviation. Solid curves represent the fit curves, obtained from the resulting fit parameters (depicted in the plot). Dashed lines depict the saturation value. Inset plot shows the data cloud with the same fit curves. b) Influence of the fit parameters on the theoretical curves. The solid black curve depicts the fit for differentiated cells (as in a) and the coloured curves are obtained by increasing or decreasing one of the two fit parameters by a factor of two. c) Theory predicts similar total numbers of microtubules for both cell states. The total number of microtubules is estimated from the spindle volume data and binned by cell volume in intervals of 500 µm^3^. The large circles depict the averages for each cell volume bin, the error bars show the standard deviation.

As all theoretical models, our model likely makes significant over-simplifications. However, our theory is in line with former quantitative descriptions of spindle scaling ^9,18,53^. Firstly, in Rieckhoff *et al*. 2020, nucleators are active in the cytoplasm and inactive when sequestered to the cell membrane, which allows for embryonic spindle scaling. In our version, we treat CPAP as the nucleator of astral microtubules, which might be direct or indirect, and here, the activity of CPAP is regulated by free tubulin, which is coupled to cell state. This hints towards a similar regulation via nucleation inhibition: in the embryonic context physical sequestration by membrane, and in the differentiation context, biochemical inhibition by tubulin where equally sized cells (with identical cell surface-to-volume ratio) show a significant difference in spindle size.

Secondly, in our model the amount of polymerised tubulin is conserved based on cell volume and we relate microtubule number to spindle volume as in Good *et al*. 2012. In the theory, the values assumed for the concentration of polymerised tubulin in the spindle bulk 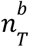 are taken from our measurements, which cannot discern which fraction of bulk tubulin is free and polymerised. However, our measurements of microtubule growth velocities and EB1 comet density within the spindle (see Fig. 2 and Suppl. Fig. 3a) imply the ratio between free and polymerised tubulin in the spindle bulk to not significantly differ between cell states.

Thus, a potential overestimate of polymerised microtubule numbers would affect both cell states and consequently would not translate into a difference in scaling behaviour.

What distinguishes our model from previous established ones ^9,18,53^ is that it proposes a regulatory mechanism for the partitioning of polymerised tubulin into either spindle bulk or astral microtubules while scaling with cell volume.

**Supplementary table S1.**
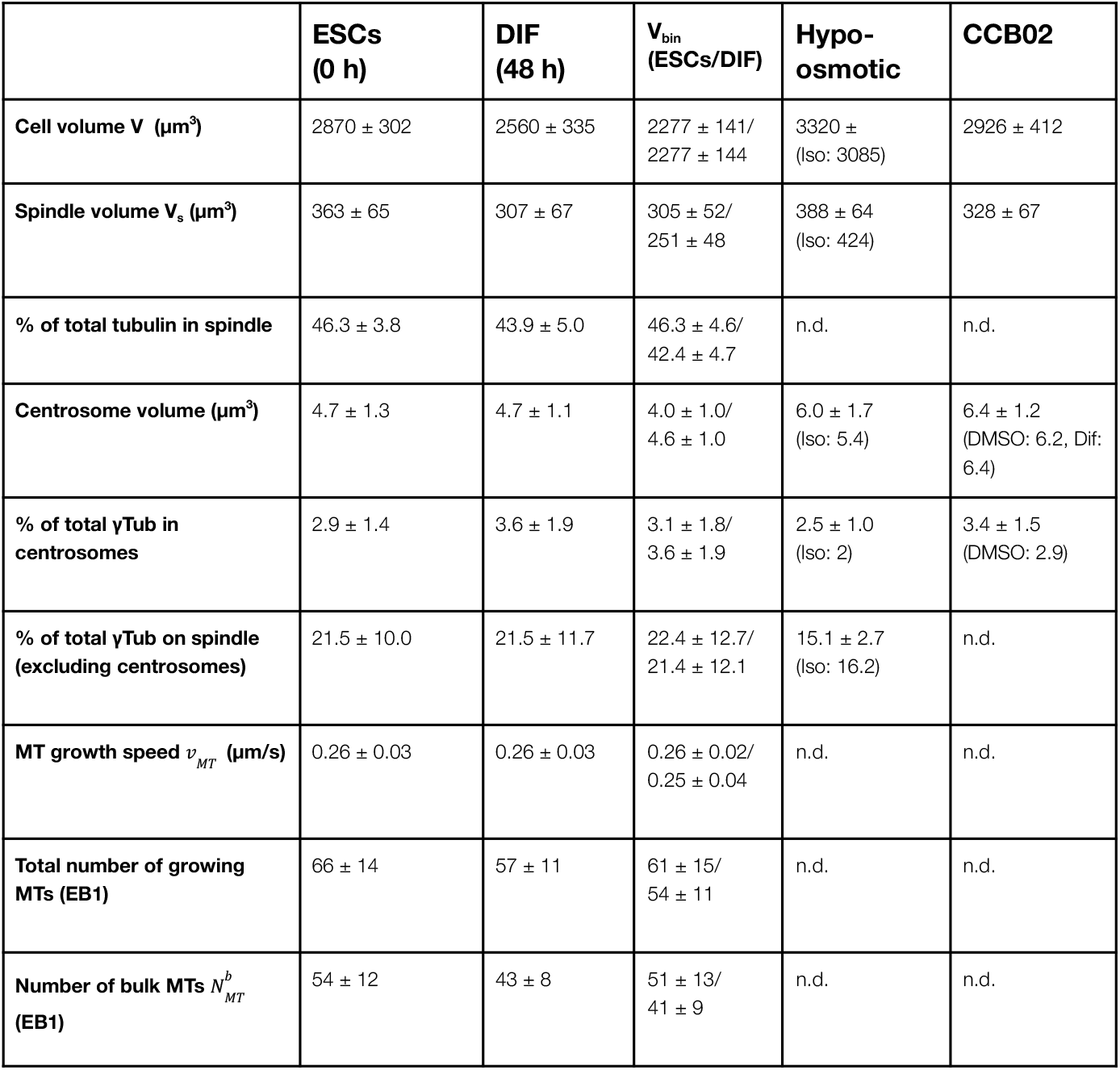
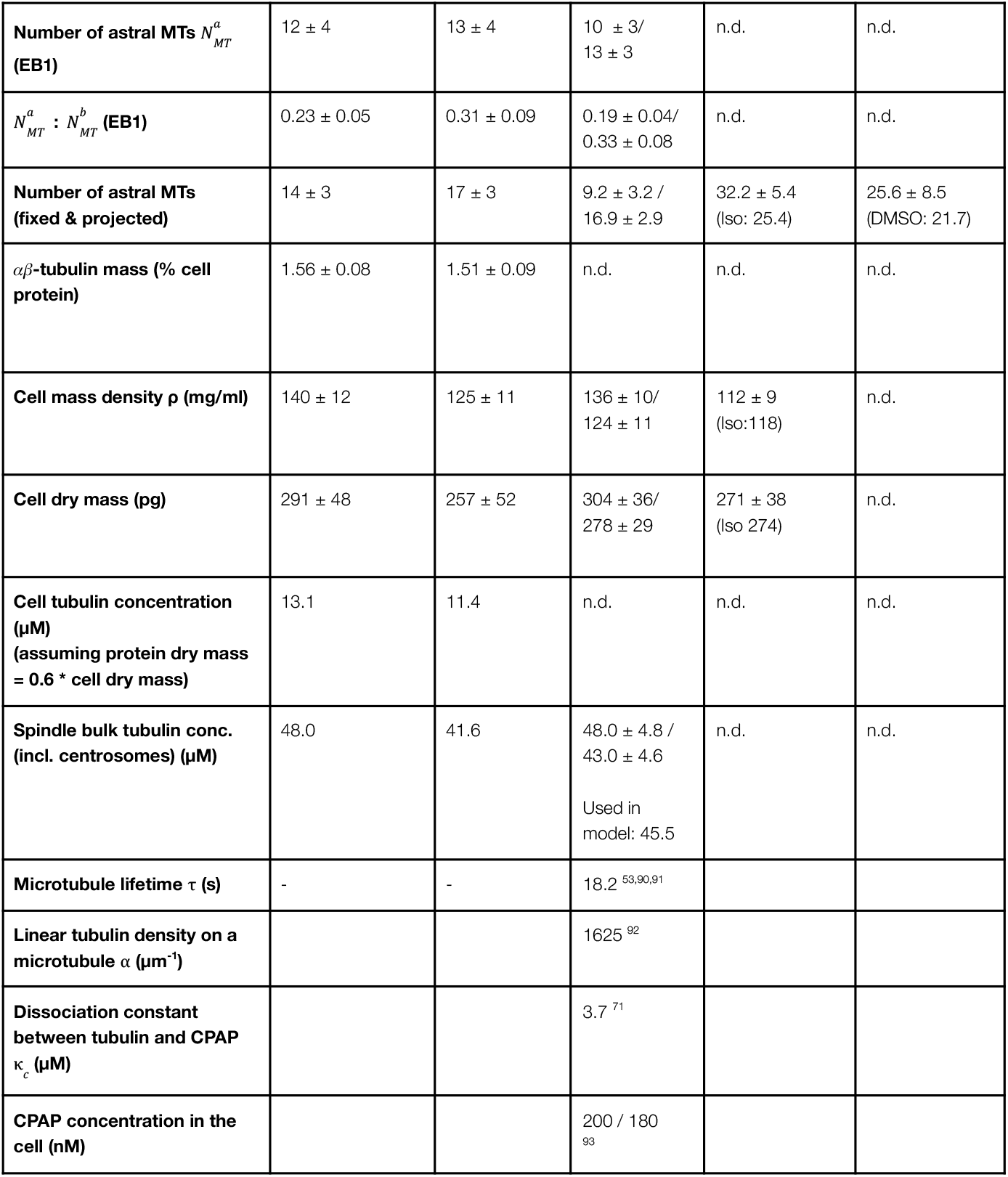
Overview summarising key measurements (mean ± standard deviation) and calculations in undifferentiated ESCs (ESCs) vs early-differentiated cells (DIF), as well as in cells during hypoosmotic treatment, or treatment with the small molecule inhibitor CCB02. V_bin_ compares ESCs vs DIF with comparable cell volumes (2000 - 2500 µm^3^). Iso: isotonic control, DMSO: control for CCB02, n.d.: not determined, γTub: γ-Tubulin.

## SUPPLEMENTARY INFORMATION

**Supplementary Figure 1:**
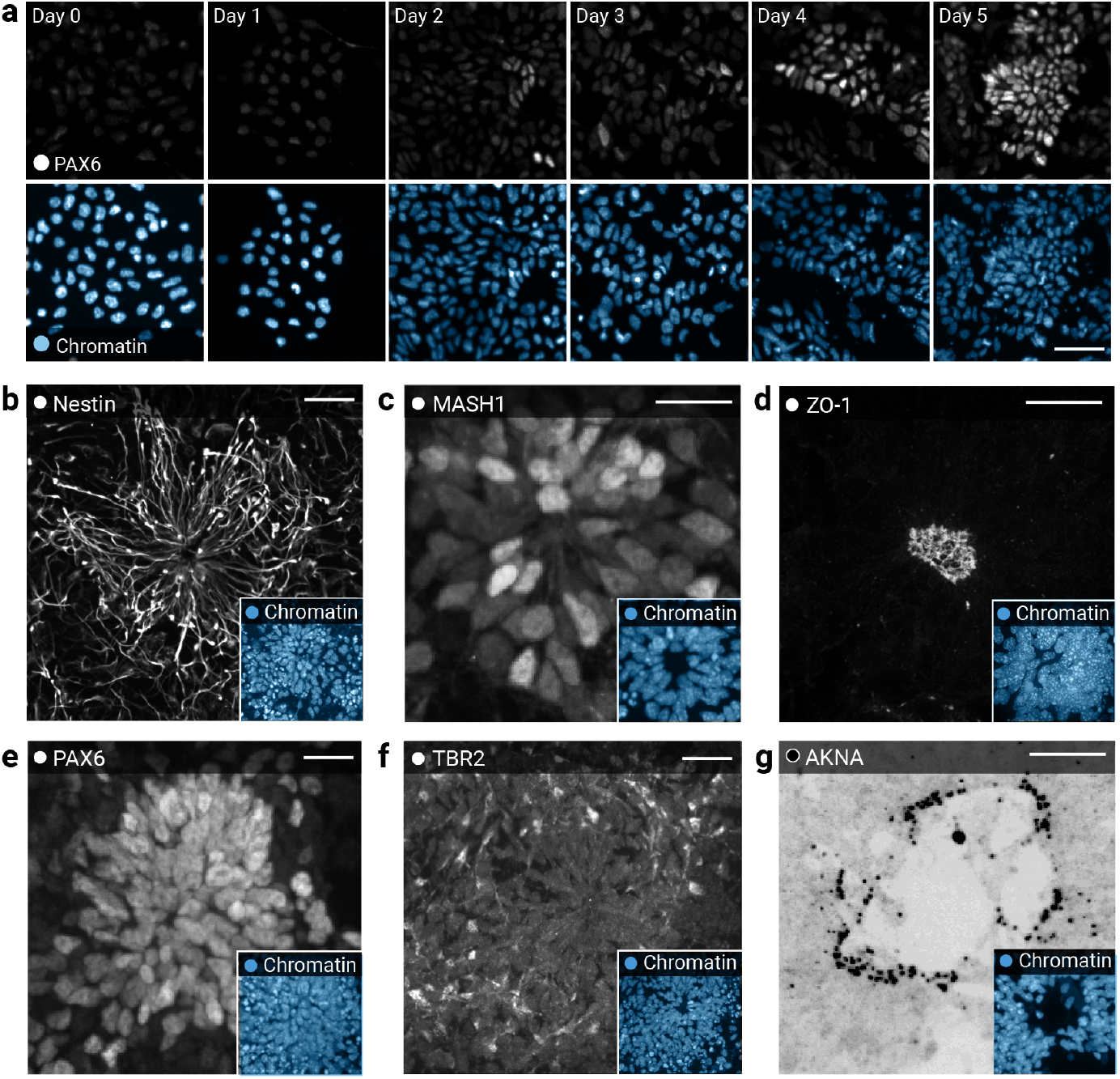
Marker expression and tissue architecture confirm successful differentiation of ESCs into neural rosettes. a) Top: Confocal data showing immunostaining of the neural marker PAX6 ^94^ in fixed ESCs (“Day 0”) and during 5 days of neural differentiation. Bottom: Nuclei stained with Hoechst. Scale bar: 50 µm. b) Immunofluorescent detection of the neural stem cell marker nestin ^95^ forming intermediate filaments in ESCs-derived rosette cells after 5 days of differentiation. Inlets show nuclei stained by Hoechst. Scale bars: 20 µm. c) As in a) but detecting MASH1 (also known as ASCL1), a proneural transcription factor ^96^, differentially expressed across rosette cells. d) As in a) but detecting ZO-1, marking tight junctions in the rosette centre (corresponding to the apical side in the mouse neuroepithelium) ^97^. e) As in a) but detecting the transcription factor PAX6, uniformly expressed in the rosette bulk indicating the presence of “apical” progenitors. f) As in a) but detecting the transcription factor TBR2 ^98^, sporadically expressed by cells in the rosette periphery, indicating “basal” or “intermediate” progenitors. g) As in a) but detecting AKNA, a centrosomal protein in neural stem and progenitor cells. AKNA^+^ centrosomes strictly localise to the “apical” pole during interphase ^99^. Lookup table of this panel is inverted in comparison with other panels for better visibility.

**Supplementary Figure 2:**
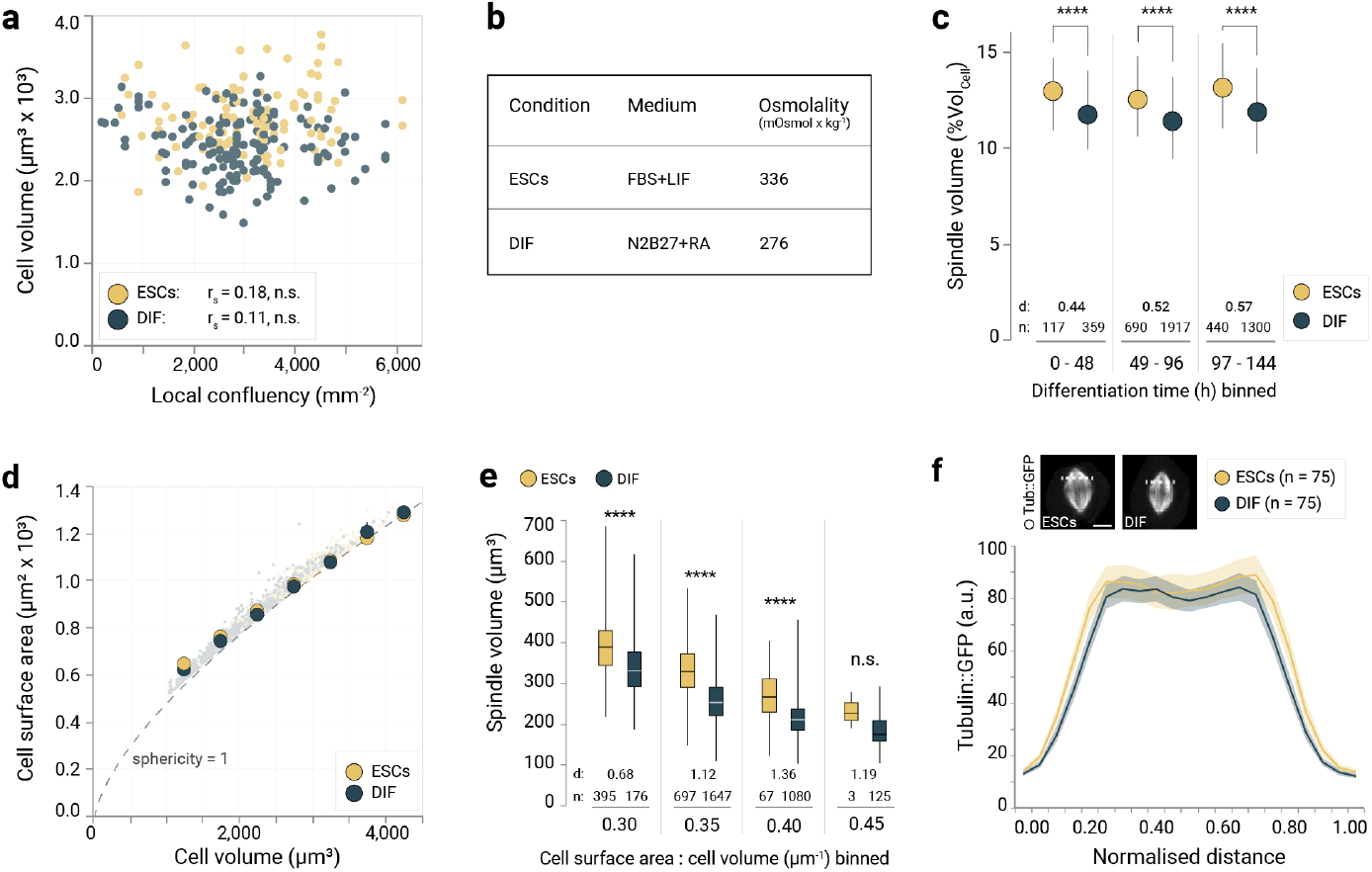
Spindle volume subscaling in differentiating embryonic stem cells (ESCs) is independent of confluency and cell geometry. a) Mitotic cell volume is independent of the local cell confluency. ESCs were incubated for 48 h either in pluripotency medium or in neural differentiation medium and then imaged live. Single data points represent individual cells (pluripotent n = 110 (yellow) and differentiation n = 151 (blue) from N = 3 experiments). r_s_: Spearman’s correlation coefficient. n.s.: not significant. b) Change of cell volume is not driven by media conditions. Osmolality of the ESCs culturing medium and of the differentiation medium. c) Cell volume occupied by the spindle in undifferentiated ESCs (yellow) and differentiating cells (blue) per differentiation time bin, showing that spindle subscaling happens from early differentiation times onwards. The circles show the means, error bars show the standard deviation. Welch’s t-test, ****: p < 0.0001. n: data points in each bin. d: Cohen’s d. d) As cell size changes, cell surface area : volume ratios (CA:CV) change. However, there is no difference between the differentiation states. Each data point represents an individual cell (ESCs n = 1088 (yellow) and differentiation n = 2921 (blue), data from N = 9 experiments). Big circles represent the median of each cell volume bin, the error bars show the interquartile range. The dotted line represents the behaviour of a perfect sphere. e) In cells with comparable CA:CV, spindle volume subscales in differentiating cells when compared to stem cells (ESCs: yellow, differentiation: blue). The boxes show the interquartile ranges, the white lines inside the boxes denote the medians, whiskers show the extrema. n: data points in each bin. d: Cohen’s d. Welch’s t-test for data in each bin, ****: p < 0.0001, n.s.: not significant. f) Spindle microtubule density stays similar before and during differentiation. Top: Maximum-projected confocal images of mitotic ESCs or early-differentiated cells (Tubulin::GFP in grey) acquired during our automated live-cell imaging protocol (Figure 1e). Bottom: Tubulin::GFP-based intensity profiles of a cross-section between spindle equator and pole in maximum-projected confocal images (top, dotted white lines). For consistency, cells were randomly drawn from a pool of cells within the 2500-3000 µm^3^ cell volume bin (ESCs n = 75 (yellow) and differentiation n = 75 (blue)). Lines denote the means, shaded areas denote the 95% confidence intervals.

**Supplementary Figure 3:**
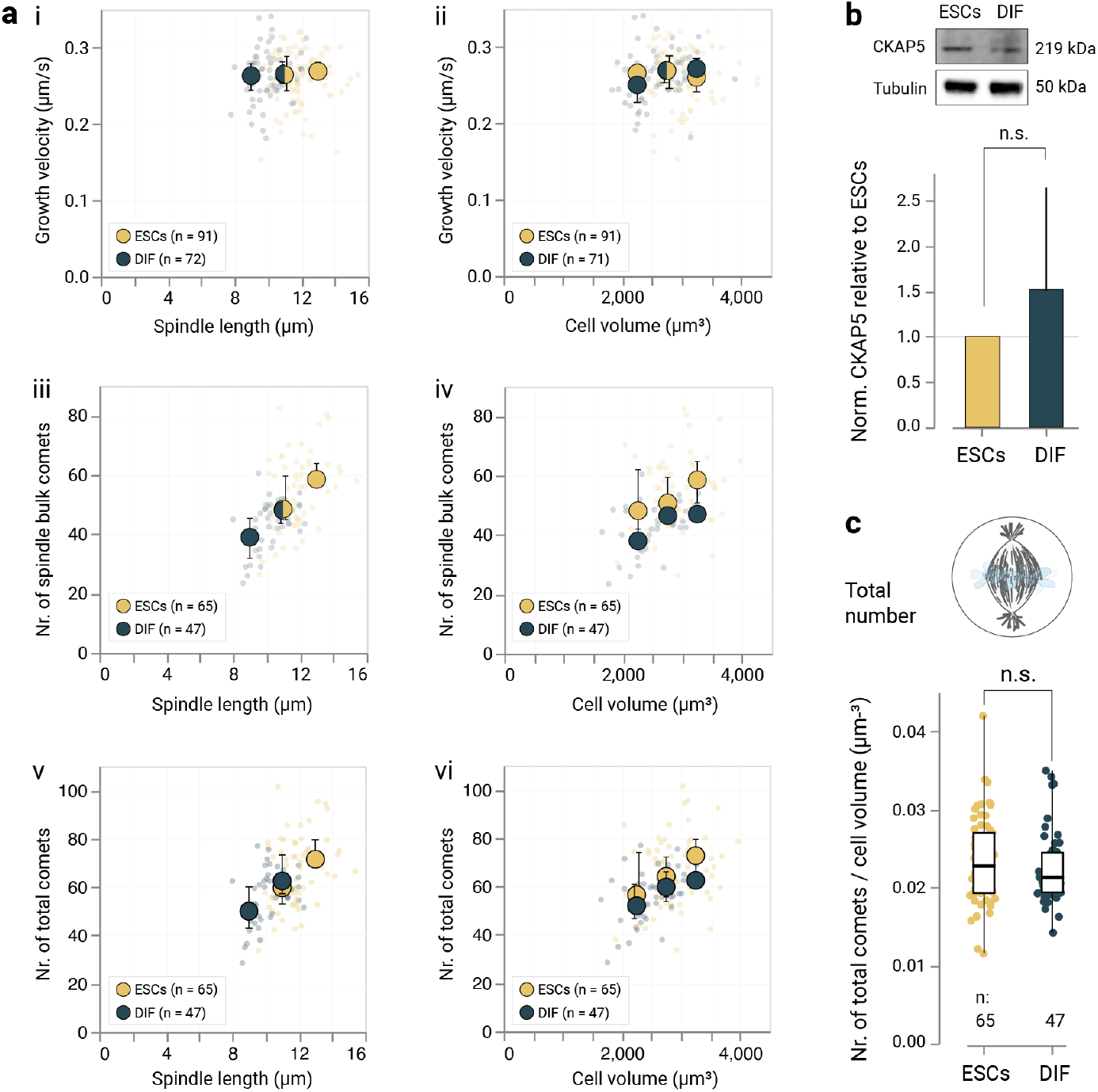
Microtubule growth speed is independent of spindle length and cell volume. a) Average single-cell microtubule growth velocities based on EB1::tdTomato comet speeds as a (i) function of spindle length and (ii) cell volume in ESCs (n = 91) or early-differentiated cells (n = 72). Average single-cell number of EB1::tdTomato spindle bulk comets as a function of (iii) spindle length and (iv) cell volume in ESCs (n = 65) or early-differentiated cells (n = 47). Average single-cell number of total EB1::tdTomato comets as a function (v) of spindle length and (vi) cell volume in ESCs (n = 65) or early-differentiated cells (n = 47). All data pooled from N = 6 experiments. Small circles show averages of individual cells, large circles represent the medians of binned data, error bars represent the interquartile ranges. b) Cellular levels of CKAP5 (also known as ch-Tog, XMAP215) after 48 h of differentiation relative to the undifferentiated ESCs, probed by western blotting and normalised to tubulin (N = 5). Bars show the mean, errors show the standard deviation. Significance tested by Welch’s t-test, n.s.: not significant. c) Number of total EB1::tdTomato comets normalised to cell volume. Each data point represents an individual cell (ESCs n = 65 and differentiation n = 47, data from N = 6 experiments). The boxes show the interquartile ranges, the black lines inside the boxes denote the medians, whiskers show the extrema. Welch’s t-test, n.s.: not significant.

**Supplementary Figure 4:**
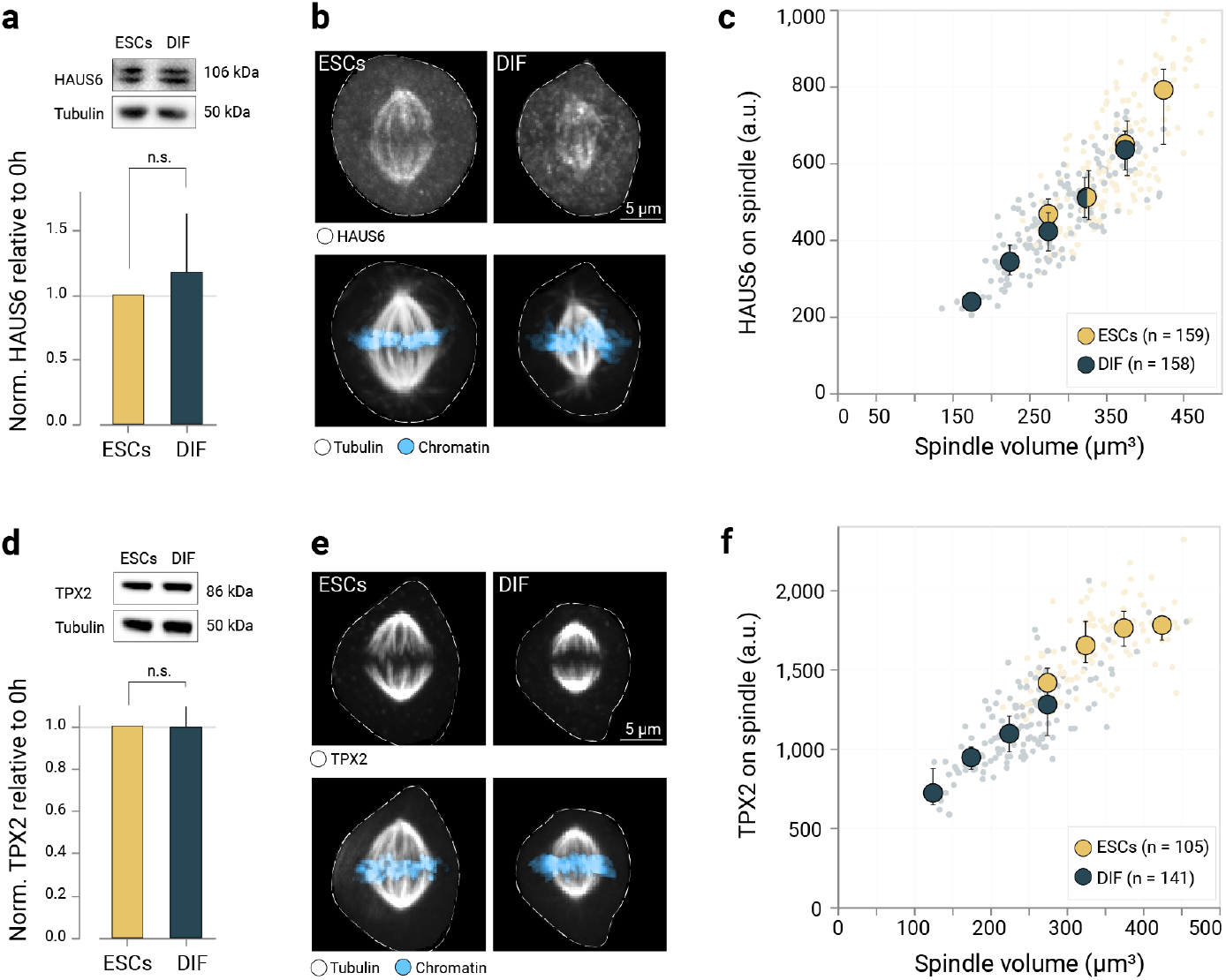
Augmin and TPX2 spindle localisation is comparable between the differentiation states. a) Cellular levels of Augmin (subunit HAUS6) after 48h of differentiation relative to the undifferentiated ESCs, probed by western blotting and normalised to tubulin (N = 3). Bars show the mean, errors show the standard deviation. Significance tested by Welch’s t-test, n.s.: not significant. b) Confocal micrographs (maximum-projected) of fixed undifferentiated ESCs or early differentiated cells at metaphase. Top: Immunostained HAUS6 signal, bottom: tubulin::GFP (grey) and chromatin counterstained by Hoechst (blue). Dotted lines indicate the cell boundaries. Scale bar: 5 µm. c) Fluorescent HAUS6 signal on spindle (normalised by total cell HAUS6 fluorescence) as a function of spindle volume. Each datapoint represents a single cell (ESCs n = 159 and differentiation n = 158, data from N = 2 experiments), the large circles denote the medians in each cell volume bin, the error bars show the interquartile ranges. d) As in a) but showing TPX2 levels. e) As in b) but showing TPX2 immunofluorescence. f) As in c) but showing TPX2 immunofluorescence (ESCs n = 105 and differentiation n = 141, data from N = 3 experiments).

**Supplementary Figure 5:**
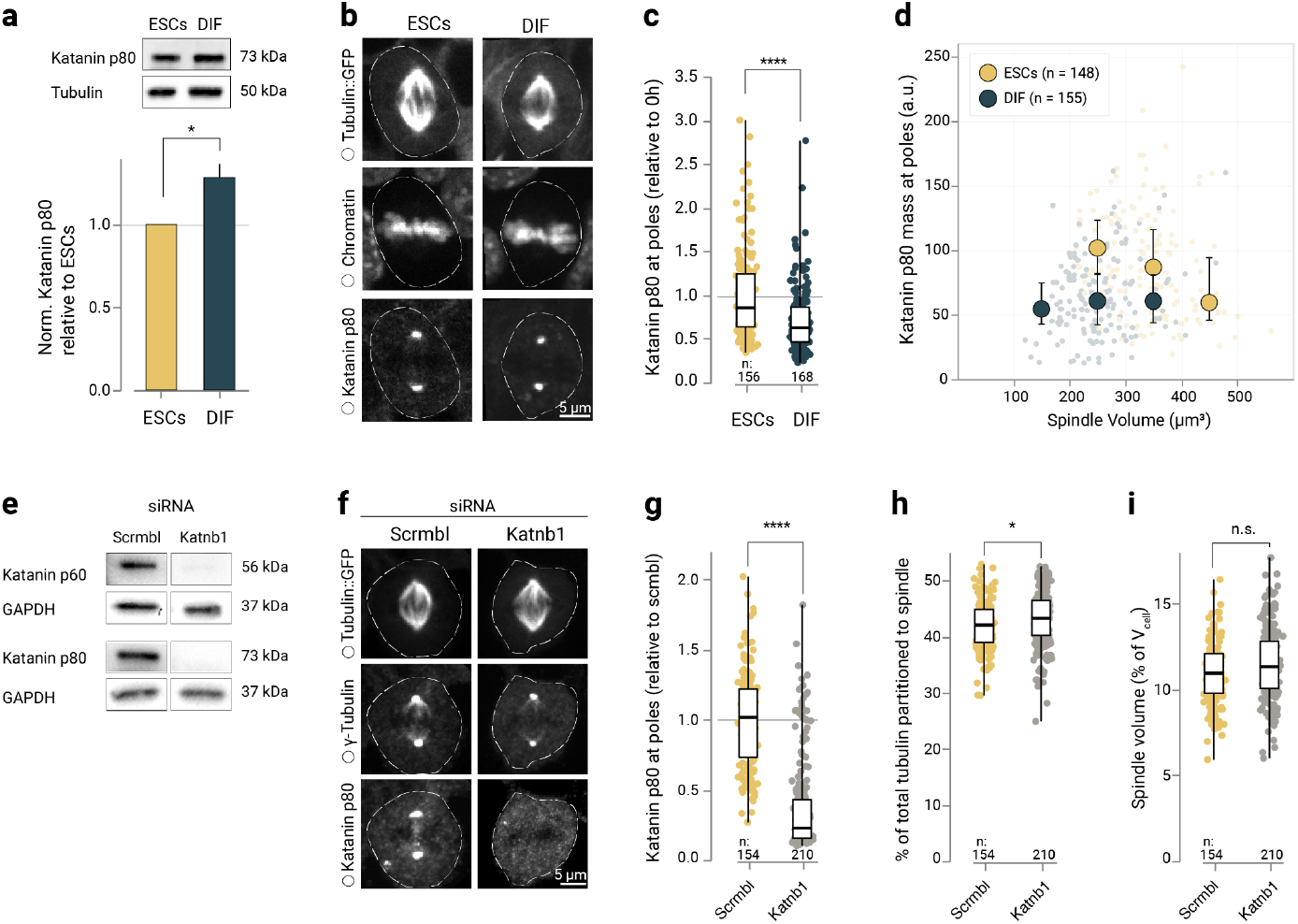
Spindle scaling in ESCs is independent of Katanin. a) Cellular levels of Katanin (subunit p80) after 48 h of differentiation relative to the undifferentiated ESCs, probed by western blotting and normalised to tubulin (N = 3). Bars show the mean, errors show the standard deviation. Significance tested by Welch’s t-test, *: p < 0.05. b) Confocal micrographs (maximum-projected) of fixed undifferentiated ESCs or early-differentiated cells at metaphase. Top: Tubulin::GFP signal, centre: chromatin counterstained by Hoechst, bottom: immunostained Katanin p80. Dotted lines indicate the cell boundaries. Scale bar: 5 µm. c) Average katanin signal at spindle poles is reduced in early-differentiating cells. Data points show individual cells (ESCs n = 156 and differentiation n = 168 from N = 3 experiments), boxes denote the interquartile ranges, horizontal lines inside the boxes denote the medians, whiskers show the extrema. Welch’s t-test,****: p < 0.0001. d) Increased Katanin p80 mass in ESCs is independent of spindle volume. Data points show individual cells (ESCs n = 148 and differentiation n = 155 from N = 3 experiments). Large circles show the medians of spindle volume bins (bin size = 100 µm^3^), error bars show the interquartile ranges. e) Representative western blots (from N = 3 experiments) showing Katanin p80 knockdown (and Katanin p60 co-depletion, as has been described previously (Jiang et al. 2017)) in ESCs using siRNAs (Katnb1) next to control using scrambled siRNAs (Scrmbl). GAPDH as loading control. f) Confocal micrographs (maximum-projected) of fixed undifferentiated ESCs after Katanin p80 knockdown (Katnb1) or control (Scrmbl). Top: Tubulin::GFP signal, centre: γ-Tubulin immunostaining as spindle pole reference, bottom: immunostained Katanin p80. Dotted lines indicate the cell boundaries. Scale bar: 5 µm. g) As in c) but showing Katnb1 siRNA-treated ESCs and control ESCs (Scrmbl) (Katnb1 n = 210 and Scrmbl n = 154 from N = 3 experiments). h) Left: Representative micrographs (maximum-projected) of 4 spindle shape phenotypes classified by severity in the siRNA experiment. Right: Percentages of the 4 spindle phenotypes within the overall population after Katnb1 siRNA treatment vs. control (Scrmbl). i) The fraction of tubulin::GFP partitioned to the spindle bulk is slightly increased upon Katanin p80 knockdown. Data points show individual cells (Katnb1 n = 210 and Scrmbl n = 154 from N = 3 experiments), boxes show interquartile ranges, horizontal bars show medians, error bars show extrema. Welch’s t-test, *: p < 0.5. j) As in i) but showing the cell volume occupied by the spindle bulk. Welch’s t-test, n.s.: not significant, p > 0.5.

**Supplementary Figure 6:**
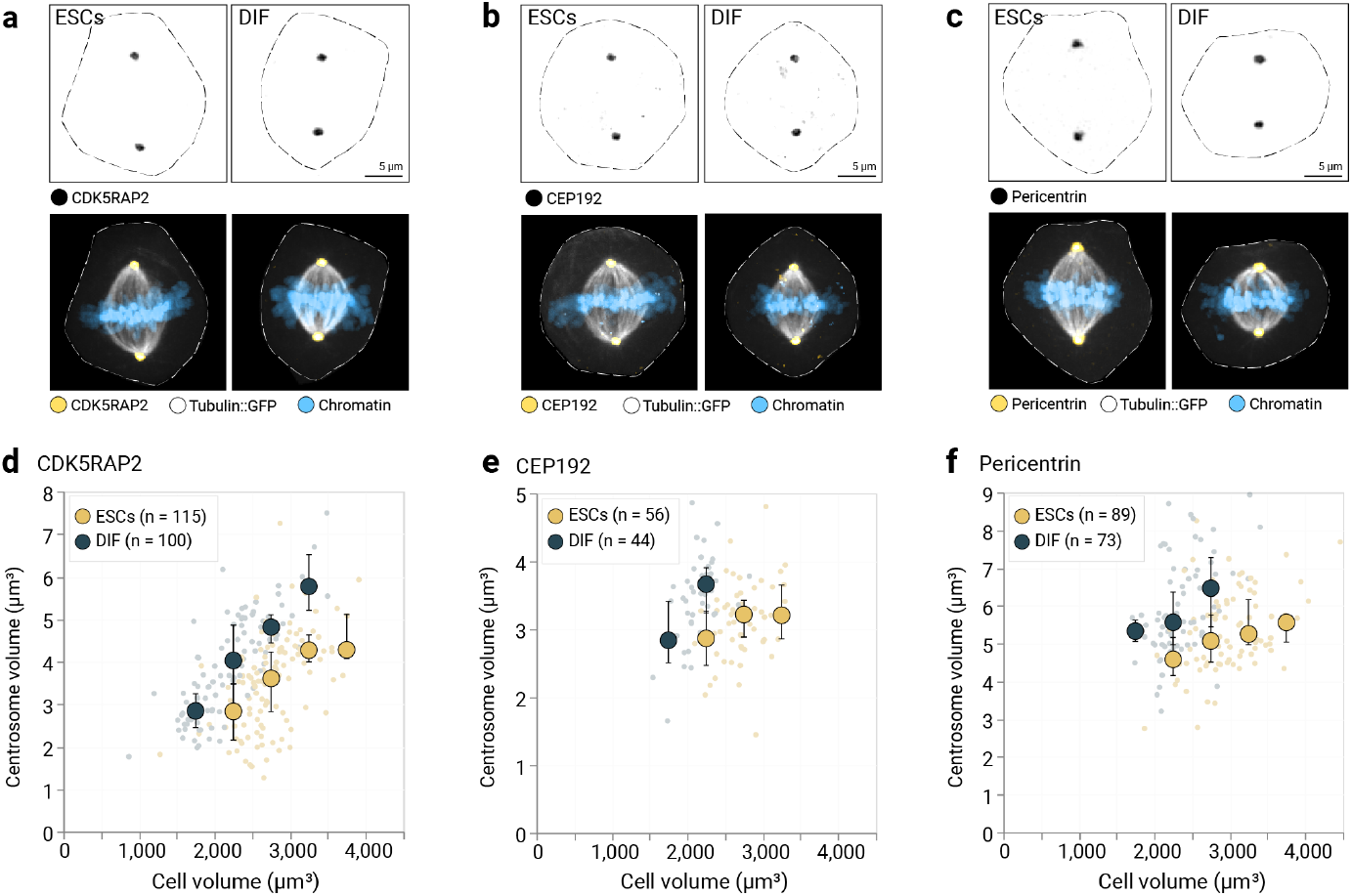
The pericentriolar material (PCM) superscales upon differentiation. a) Confocal micrographs (maximum-projected) of fixed undifferentiated ESCs or early differentiated cells at metaphase. Top: Immunostained CDK5RAP2 signal (inverted grayscale), bottom: immunostained CDK5RAP2 (yellow), tubulin::GFP (grey) and chromatin counterstained by Hoechst. Dotted lines indicate cell boundaries. Scale bar: 5 µm. b) As in a) but using CEP192 as PCM marker. c) As in a) but using Pericentrin as PCM marker. d) The PCM is enlarged in early-differentiating cells. Scaling relationship between cell volume and centrosome volume based on CDK5RAP2 immunofluorescence. Each data point represents a single cell (ESCs n = 115 cells, differentiation n = 100 cells from N = 3 experiments), the large circles denote the medians in each cell volume bin, the error bars show the interquartile ranges. e) As in d) but based on CEP192 immunofluorescence (ESCs n = 56 cells, differentiation n = 44 cells from N = 1 experiment). f) As in d) but based on Pericentrin immunofluorescence (ESCs n = 89 cells, differentiation n = 73 cells from N = 3 experiments).

**Supplementary Figure 7:**
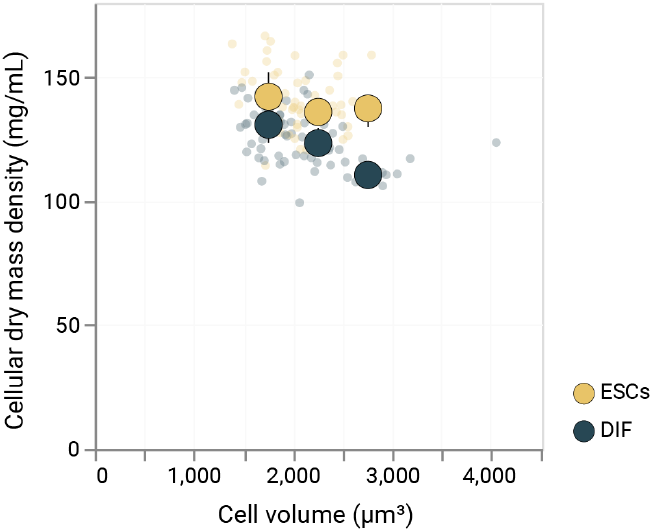
The cytoplasm in differentiating cells is diluted across all cell sizes. Scaling relationship between cell volume and cellular dry mass density. Each data point represents a single cell (ESCs n = 61 cells, differentiation n = 71 cells from N = 3 experiments), the large circles denote the medians in each cell volume bin, the error bars show the interquartile ranges.

**Supplementary Movie 1: Emergence of neural rosettes in adherent ESC differentiation culture**.

Confocal time-lapse recording of ESCs expressing tubulin::GFP (grey) undergoing neural differentiation, chromatin stained using SiR-DNA (blue). In later time points, cells self-organise to form rosette-like tissue architectures. Scale bar: 25 µm, time stamps are hh:mm (15 min per frame, total of 112.25 hours).

**Supplementary Movie 2: Interkinetic nuclear migration in ESC-derived neural rosettes**. Confocal time-lapse recording of rosette cells (tubulin::GFP in grey, chromatin stained using SiR-DNA, blue). Mitoses of rosette cells exclusively take place in the centre of the rosette (corresponding to the “apical” side), whereas the nuclei migrate to the periphery of the rosette, a phenomenon called interkinetic nuclear migration. Scale bar: 25 µm, time stamps are hh:mm (5 min per frame).

**Supplementary Movie 3: Visualising growing microtubules in live mitotic cells**.

Confocal time-lapse recording (single z plane) of a mitotic undifferentiated embryonic stem cell expressing EB1::tdTomato (grey). EB1 labels growing microtubules plus ends. Scale bar: 5 µm, 0.4 seconds per frame.

**Supplementary Movie 4: Increased EB1 localisation at the centrosomes upon differentiation**.

Confocal time-lapse recordings (single z planes) of mitotic early-differentiated cell (left) and an undifferentiated embryonic stem cell (right) expressing EB1::tdTomato (grey). The EB1 signals are cumulative per frame. The global histograms of the individual movies are normalised to the last frame. Scale bar: 5 µm, 0.4 seconds per frame.

## METHODS

### Quantification and statistical analysis

In each figure legend, details about the quantifications are provided, including the number of analysed cells and spindles measured (n) and the effect size (Cohen’s d). The effect size was calculated by the following formula: d = (mean_of_ESCs - mean_of_DIFs)/s where s is the pooled standard deviation of the parameter.

Data analysis was performed using the *pandas* library ^100^ and the *NumPy* library ^101^ in *Jupyter Notebooks* (using Python 3.7.1). Statistical tests were performed using the *SciPy* library (Welch’s t-test for unequal variances, Wilcoxon signed-rank test) ^102^ or the *Statsmodels* package (ANOVA) ^103^. Spearman’s correlation coefficients were determined using the *SciPy* stats.spearman package. The P values given show the probability of an uncorrelated system producing data sets that have a correlation at least as extreme. Minimum-maximum normalisation (EB1 half-spindle intensity profiles) was performed using the *Scikit-learn* library ^104^.

### Stem cell culture and differentiation

R1/E mouse embryonic stem cells (ESCs) stably transfected with bacterial artificial chromosomes harbouring the eGFP-fused coding region of human β5-tubulin and its regulatory sequences for native expression levels (gift from the Hyman lab, Dresden) ^105^ were routinely cultured in FBS+LIF medium (16% FBS (gibco), non-essential amino acids (gibco), 50 µM β-mercaptoethanol, pen/strep (Invitrogen), 10^3^ units/mL recombinant mouse LIF (Sigma-Aldrich) in high-glucose DMEM supplemented with L-Gln, Pyruvate). After thawing from N_2_ storage, ESCs were passaged routinely every 48 h. Tissue culture dishes (100 × 25 mm) were freshly coated with 0.1% gelatin (in ddH_2_O). The cells were washed with pre-warmed 1x PBS (pH 7.4) and detached from the vessel (pre-warmed Accutase solution (Invitrogen) at RT). Detachment was stopped through the addition of warm FBS+LIF medium. Cells were centrifuged for 2 min at 200 x g. Finally, cells were transferred to the gelatin-coated dish containing pre-warmed FBS+LIF medium at a density of 25,000 cells per cm^2^. The culture dish was incubated at 37 °C, 95% humidity and 5% CO_2_.

Alternatively, ESCs were cultured using 2i+LIF medium (N2 in DMEM/F12 (gibco) mixed 50:50 with B27 (200x, w/o vitamin A, gibco) in neurobasal medium (gibco), supplemented with L-Gln and 50 µM β-mercaptoethanol, pen/strep (Invitrogen), 3 µM CHIR99021 and 1µM PD0325901) prepared from N2 supplement (100x) made in-house (0.6 mg/mL progesterone (Sigma-Aldrich), 1.6 mg/mL putrescine (Sigma-Aldrich), 10 mg/mL apo-transferrin (Sigma-Aldrich), 3 µM sodium selenite (Sigma-Aldrich), 2.5 mg/mL insulin (Sigma-Aldrich), 0.5% bovine albumin fraction V (gibco) in DMEM/F12 (gibco)). The cells were passaged as described above but using 2i+LIF instead of FBS+LIF and at a reduced cell seeding density of 15,000 cells per cm^2^.

For neural differentiation, undifferentiated ESCs were taken up in N2B27 medium (N2 (see above) in DMEM/F12 (gibco) mixed 50:50 with B27 (200x, plus vitamin A, gibco) in neurobasal medium (gibco), supplemented with L-Gln and 50 µM β-mercaptoethanol, pen/strep (Invitrogen), 500 nM retinoic acid (Roth)) and seeded onto laminin-511 (L511, 2.5 µg/mL in 1x PBS (containing Ca^2+^ and Mg^2+^)) coated dishes or imaging wells at a density of 19,000 cells per cm^2^ and incubated at 37 °C, 95% humidity and 5% CO_2_. For differentiation times longer than 48 h, cells were passaged after 48 h and replated at a density of 200,000 cells per cm^2^.

Cells were routinely tested for Mycoplasma contamination using a commercial detection kit.

### RNAi

For siRNA transfection, ESCs were seeded at a seeding number of 3,500 cells per cm^2^ on L511-coated dishes. After 24 h, the cells were transfected using the Lipofectamine 3000 kit (Thermo) and 20 nM of siRNAs (katnb1 5’-GGACUACAGGAGAUAUCAAtt-3’ or scrambled 5’-UUCUCCGAACGUGUCACGUtt-3’) according to the manufacturer’s instructions. After 48 h of knockdown, the cells were either fixed, immunostained and imaged as described in Section “Pericentriolar material”, or lysed and prepared for western blotting as described in Section “Western blotting”.

### Live-cell confocal microscopy

For single time-point imaging of mitotic cells, cells were seeded and differentiated as described above in L511-coated imaging plates (polymer bottom, Ibidi). Cells were imaged using a Nikon spinning disk confocal equipped with an incubation chamber (set to 37 °C, 95% humidity and 5% CO_2_), a Andor Revolution SD System (CSU-X), an iXon3 DU-888 Ultra EMCCD camera and a 60x Plan Apo oil objective (numerical aperture = 1.40).

Fields of view were selected manually and confocal series (z range of 23 µm at a step size of 0.3 µm) were recorded using the triggered acquisition function, using a multi-wavelength filter and the 488 nm and 640 nm laser lines at a power of 5% laser power and 200 ms exposure time.

### Adaptive feedback microscopy

The samples were prepared using glass-bottom 4-well imaging slides (Ibidi), coated with L511 (2.5 µg/mL in 1x PBS (containing Ca^2+^ and Mg^2+^)). Five hours before the imaging, cells were passaged as described above. Because undifferentiated ESCs would at some point become too confluent, the ESCs were seeded at an initial population density of 2,000 cells per cm^2^. Differentiating ESCs were seeded at the usual initial population density of 19,000 cells per cm^2^ to drive differentiation efficiently. The data sets were recorded on two laser-scanning confocal imaging systems, the Zeiss LSM 800 and LSM 900, equipped with custom-built, temperature, CO_2_ and humidity controlled incubation chambers (Electronic and mechanical workshops, EMBL Heidelberg). A C-Apochromat 40x water objective (numerical aperture = 1.2) was used. Signals were detected using GaAsP photomultiplier tubes. To sustain water immersion of the objective during continuous imaging over days, the objective was equipped with a custom-built automated water immersion system (Electronic and mechanical workshops, EMBL Heidelberg). The pump was automatically triggered by the adaptive feedback microscopy pipeline (see below) to replace the water in 20-minute intervals when the microscope was not imaging. To counteract evaporation and re-feed the cells in the culture wells, the media were carefully replaced by hand every 24 h.

We set up and used the adaptive feedback microscopy pipeline to automatically locate mitotic cells within a larger population of cells over days and image them in 3D with high resolution (see Figures 1e, f). The pipeline consists of 4 image acquisition jobs and 3 image analysis jobs as described below. The pipeline was driven by Zeiss *ZEN Blue* macros, controlling the acquisition process, and an analysis protocol set up in *FIJI* ^*106*^ using the *AutoMicTools* package.

1. Low-zoom autofocus. After moving the stage to the next predefined position in each imaging cycle, the focal plane was re-determined by using a reflection signal from the top surface of the coverslip. A fast XZ scan was performed using the piezo drive and the reflection signal was calculated by online image analysis and fed back to the microscope to trigger the acquisition of the low-zoom population image in focus. Settings: see Methods Table 1.
2. Low-zoom population image. Two-channel (DNA, GFP) overview images of cells at predefined positions in the wells with non-differentiating / differentiating cells were acquired by recording 2×2 tile scans with a 10% overlap and immediately stitched in the *ZEN Blue* software. Settings: see Methods Table 1. The stitched image was processed to identify positions of mitotic cells.
3. High-zoom autofocus. After receiving the xy coordinates of the positive hits, the pipeline determined the central focal plane for each mitotic cell for the subsequent high-content acquisition (4). For that purpose a stack at low lateral resolution, high zoom factor and extended z-range was automatically acquired for each cell in question. The online image analysis in *AutoMicTools* identified the mask of the tubulin::GFP-labelled spindle and found its brightest z section to correct lateral and axial position of a high resolution acquisition (see step 4). Settings: see Methods Table 1.
4. High-zoom single-cell image. Having received the final xzy coordinates, the positive hits were recorded in 2 channels with high spatial resolution. Settings: see Methods Table 1.

For one imaging cycle (interval of 10 min), the microscope moved to the first predefined xy position and recorded the low-zoom autofocus image (1) that was analysed using the *AutoMicTools* plugin in *FIJI*. The correct focal z plane was sent back to the microscope that subsequently performed job (2). In the dedicated *AutoMicTools* online image analysis job the GFP channel in the overview raw image was first maximum-projected. Then positions of mitotic cells were identified using the *Multi-Template Matching* plugin ^107^where we provided a template for object detection of tubulin::GFP-labelled spindles from our previous data. We limited the number of positive hits to a maximum of 8. The list with the xy coordinates was fed back to the microscope to proceed with jobs (3) and (4) for each entry. Once completed, the cycle started anew at the next predefined position.

**Methods Table 1:** Adaptive feedback microscopy settings.

|  | 1. Low-zoom autofocus | 2. Low-zoom population image | 3. High-zoom autofocus | 4. High-zoom single-cell image |
| --- | --- | --- | --- | --- |
| Voxel width | 4 $\mu\text{m}$ | 0.312 $\mu\text{m}$ | 0.71 $\mu\text{m}$ | 0.228 $\mu\text{m}$ |
| z-dimension | 0.1 $\mu\text{m}$ | 3 $\mu\text{m}$ | 0.5 $\mu\text{m}$ | 0.75 $\mu\text{m}$ |
| Nr. of z | 300 | 3 | 61 | 32 |
| xy dimensions | 160 $\mu\text{m}$ x 4 $\mu\text{m}$ | 607 $\mu\text{m}$ x 607 $\mu\text{m}$ | 22.8 $\mu\text{m}$ x 22.8 $\mu\text{m}$ | 22.8 $\mu\text{m}$ x 22.8 $\mu\text{m}$ |
| Pinhole size | 41.5 $\mu\text{m}$ | 218.4 $\mu\text{m}$ | 42.2 $\mu\text{m}$ | 49.3 $\mu\text{m}$ |
| Scan speed / Averaging | 8 (bi) / 1 | 8 (uni) / 4 (mean) | 16 (uni) / 1 | 10 (uni) / 2 (mean) |
| Pixel dwell time | 2.18 $\mu\text{s}$ | 4.89 $\mu\text{s}$ | 0.76 $\mu\text{s}$ | 3.39 $\mu\text{s}$ |
| Magnification | 1 | 0.5 | 7.7 | 7.7 |
| Laser lines (power) | 640 nm (5%) | 488 nm (10%), 640 (5%) | 488 nm (10%) | 488 nm (10%), 640 (5%) |
| Detection | 630-700 nm | 410-559 nm, 656-700 nm | 410-559 nm | 410-559 nm, 656-700 nm |

### Quantification of cell cycle duration

Cell families were traced using the manual mode in *TrackMate* ^108^ in the concatenated, max-projected and median(sigma=2)-filtered time-lapse movies generated via adaptive feedback microscopy. The time interval between two track division events was read out as the cell cycle duration.

### Quantification of cell and spindle morphology

Cell volumes and cell surface areas were calculated via *MorphoLibJ* ^109^ in *FIJI* after segmenting cells in 3D as described in Kletter et al. 2021, or using the *Volume Manager* plugin within the plugin package provided by the Scientific Computing Facility of the MPI-CBG (Dresden, https://sites.imagej.net/SCF-MPI-CBG/). Spindle volumes were determined using *Spindle3D* (0.80) ^51^ after excluding all voxels outside the cell binary masks. For clarity, microscopic images shown in the figures only show voxels within the cell binary masks.

### Quantification of microtubule growth velocity and EB1 comet numbers

Undifferentiated ESCs were transiently transfected following manufacturer’s instructions (Lipofectamine 3000, Thermo) with plasmids carrying the EB1::tdTomato coding sequence under a CMV promoter (Addgene 50825). After overnight transfection, the cells were plated onto L511-coated wells of a 24-well polymer bottom imaging plate (Ibidi) at a seeding density of 19,000 cells per cm^2^, and either maintained in FBS+LIF medium or in N2B27 medium for neural differentiation.

After 48 h, cells were imaged on a Nikon spinning disk confocal equipped with an incubation chamber (set to 37 °C, 95% humidity and 5% CO2), a Andor Revolution SD System (CSU-X), an iXon3 DU-888 Ultra EMCCD camera and a 100x Plan Apo oil objective (numerical aperture = 1.45). A confocal series of the tubulin::GFP signal was recorded (step size of 0.75 μm), using a multi-band filter and the 488 nm laser at 5% power and 200 ms of excitation duration. To capture fluorescent EB1 comets, after selecting the most central position within the spindle, a single-plane time-lapse recording (400 ms frames for 1 min) was recorded using a single-band filter in combination with the 561 nm laser at 5% and 300 ms of excitation duration. Spindle lengths were manually determined in *FIJI* by calculating the 3D vector length between the two spindle poles as seen in the tubulin::GFP signal. Cells were segmented and volumetrically quantified as described above.

For automated single-particle tracking using the *FIJI* plugin *TrackMate* ^108^, the recordings were processed using a median filter (sigma = 1 μm) and a rolling-ball background subtraction (radius of 1 μm). In *TrackMate*, particles were detected using the LoG (Laplacian of Gaussian) algorithm (sub-pixel localisation, estimated blob diameter 0.5 μm, no median filter, threshold (100-130) set manually depending on the cell-specific EB1::tdTomato signal). For the generation of tracks, the LAP tracker algorithm was used with the following settings: frame linking max distance 0.3 μm, gap closing not allowed, track splitting not allowed, track merging not allowed. The results table containing the track velocities was exported. Data integration and refinement was performed using the *pandas* library and *Jupyter Notebooks* (Python 3.7.1). Tracks were filtered by duration (2-4 s), median track quality (200-600) and track start and end within the movie (Frames 0-30).

Comet numbers were determined manually. For each recording, the numbers of spindle bulk EB1 comets and astral EB1 comets (amounting to the total number of EB1 comets) were determined in frame number 1, frame number 20 and frame number 40 and averaged. Lastly, the EB1::tdTomato signals were sum-projected in *FIJI*. To minimise the effect of spindle rotations during the recordings, only the first 20 seconds were considered. Again in *FIJI*, using the line tool at an averaging of 15 pixels, half-spindle profiles were drawn starting from one pole to the spindle equator. Projections were only included if the centre of the spindle and at least one pole was clearly in focus. The profiles were assembled in *Jupyter Notebooks* (Python 3.7.1). For each profile, the x-axis (corresponding to the half-spindle distances) and the y-axis (corresponding to the summed up EB1 signal) were minimum-maximum normalised using the min_max_scaler.fit_transform function in the *Scikit-learn* library ^104^.

### Pericentriolar material

Cells were seeded onto L511-coated 24-well imaging plates (Ibidi) for embryonic stem cell culture or differentiation as described above. After 48h of incubation, cells were chemically fixed using 3.2% PFA (in 1x PBS) for 10 min at 37 °C. After three washes with 1x PBS and 10 min quenching with 0.1 M glycine at RT, the cells were treated with BSA blocking buffer (3% BSA/0.5% TX100 in 1x PBS) for 1 h at RT. Primary antibody incubation was performed in BSA blocking buffer for 1 h at RT. Because of overlapping host origins, primary stainings were performed separately (CDK5RAP5 (Sigma 06-1398), 1:500; CEP192 (Proteintech 18832-1-AP), 1:500; Pericentrin (Abcam 4448), 1:500; γ-Tubulin (Sigma T6557), 1:100). Secondary antibody incubation was performed for 1 h at RT using anti-mouse or anti-rabbit AlexaFluor647 secondary antibodies diluted 1:000 in BSA blocking buffer.

The fixed and stained cells were imaged on a Nikon spinning disk confocal microscope at room temperature, equipped with an Andor Revolution SD System (CSU-X), an iXon3 DU-888 Ultra EMCCD camera and a 100x Plan Apo oil objective (numerical aperture = 1.45) or a 60x Plan Apo oil objective (numerical aperture = 1.40). Three-channel confocal series (chromatin, tubulin::GFP, antibody-staining) were recorded using a z-step size of 0.3 µm. Using single-band filters, stacks of the individual channels were recorded serially (405 nm: 5% laser power, 488 nm: 18% laser power, 640 nm: 5% laser power). Excitation duration per slice was 200 ms.

In *FIJI*, metaphase cells were first selected and cropped from the raw microscopy files. Next, the cells were 3D segmented as described above, and non-cell voxels were eliminated. The spindle segmentation was performed using the tubulin::GFP signals in *Spindle3D*.

For 3D segmentation of the PCM the PCM signal was thresholded using the maximum entropy algorithm in *FIJI*, considering the histogram from all optical sections. This method was robust because all non-cell voxels were already eliminated from the confocal stack, and so the thresholding only saw voxels belonging to either the PCM, or to cell regions outside the PCM. This thresholding step yielded the centrosome binary mask (containing both centrosomes). Volumes of the cells were calculated from the cell segmentation binary masks, and the volumes of the centrosomes were calculated from the centrosome binary mask using the Analyze Regions 3D function in *MorphoLibJ* ^109^. The average cellular fluorescence of the channel containing the PCM signal was determined using the *FIJI 3D Suite* ^110^ and the cell binary mask. This raw cell average was background-corrected by subtracting the minimum voxel value within the cell binary mask, robustly representing the system-specific noise floor. Analogously, this was repeated for the average signal in the centrosome binary mask. To quantify the average fluorescent γ-Tubulin signal residing in the spindle, the binary spindle mask from the *Spindle3D* segmentation was applied on the PCM channel using the *FIJI 3D Suite*. The average signals from the centrosome mask and the spindle mask, respectively, were normalised using the cell average signal.

### Quantification of astral microtubules

For this work, we define astral microtubules as the subpopulation of mitotic microtubules that originate at the centrosomes and are not incorporated into the spindle bulk. To visualise astral microtubules, cells were fixed and stained with anti-tubulin antibodies. Cells were seeded onto L511-coated 24-well imaging plates (Ibidi) for embryonic stem cell culture or differentiation as described above. After 48 h of incubation, cells were chemically fixed for 10 min at 37 °C with tubulin fixation buffer (3.2% paraformaldehyde / 0.1% glutaraldehyde in 1x PBS) prewarmed to 37 °C. The cells were washed three times with 1x PBS. For quenching, the cells were first treated with 0.1% NaBH4 in 1x PBS for 7 min at RT and for 10 min with 100 mM glycine in 1x PBS. Blocking was performed using BSA blocking buffer (3% BSA / 0.5% TX-100 in 1x PBS) for 1 h at RT. The cells were incubated for 1 h at RT with BSA blocking buffer supplemented with anti-alpha-tubulin monoclonal antibodies (DM1a (Sigma T-6199, mouse origin), diluted 1:1000 and Yol1/34 (Bio-rad MCA78G, rat origin), diluted 1:000). After three washes with 1x PBS, cells were incubated with secondary antibodies (anti-mouse AlexaFluor647 (Thermo) 1:000 in blocking buffer and anti-rat AlexaFluor647 (Thermo) 1:1000 in blocking buffer). After staining the DNA with Hoechst 33342 (1:10,000 in 1x PBS) for 5 min at RT, cells were again washed three times with 1x PBS.

The fixed and stained cells were imaged on a Nikon spinning disk confocal microscope at RT, equipped with an Andor Revolution SD System (CSU-X), an iXon3 DU-888 Ultra EMCCD camera and a 100x Plan Apo oil objective (numerical aperture = 1.45). Three-channel confocal series (chromatin, tubulin::GFP, antibody-staining) were recorded using a z-step size of 0.3 µm. Using single-band filters, stacks of the individual channels were recorded serially (405 nm: 5% power, 488 nm: 18% power, 640 nm: 5% power). Excitation duration per slice was 200 ms.

The cells and spindles were 3D segmented and morphometrically analysed as described above using the tubulin::GFP signals. The stained microtubule signals were maximum projected in *FIJI*. The individual astral microtubules were traced using the free-hand selection tool. The single-cell tables containing numbers and lengths of astral microtubules were concatenated in a *Jupyter Notebook* (Python 3.7.1) and merged with the quality-controlled cell and spindle morphometric data.

### Quantitative tubulin western blotting

Whole cell lysates were prepared from adherent ESCs throughout differentiation. The lysis of cells from different stages (24 h – 120 h) of differentiation was synchronised by staggering the differentiation onset. Using the differentiation procedures as described above, cells were seeded onto L511-coated 6-well plates, 3 replica wells per differentiation day. For the “0 h” time point, the ESCs were seeded at a density of 19,000 cells per cm^2^ and incubated in FBS+LIF for 48 h. Before lysis, cells were washed twice with room temperature 1x PBS.

Plates were placed on ice and cells were incubated with cold, protease inhibitor-supplemented RIPA lysis buffer (150 mM NaCl / 50 mM Tris-HCl (pH 8.0) / 1% Nonidet P-40 / 0.5% sodium deoxycholate / 0.1% SDS in ddH_2_O) and scraped off using a cell scraper. Lysates were collected and incubated for 15 min on ice. The lysates were sonicated in a water bath sonicator three times for two seconds each, with 1 min recovery periods on ice between the pulses. The lysates were incubated another 15 min on ice before centrifuging at 13,000 x g for 5 min at 4 °C. The supernatants were taken off and protein concentrations were determined using a bicinchoninic acid (BCA) assay (Thermo), following the manufacturer’s instructions.

Whole-cell RIPA lysates were diluted 1:4 with SDS sample buffer (4x) (200 mM Tris-Cl (pH 6.8), 400 mM DTT, 8% SDS, 0.4% bromophenol blue, 40% glycerol), briefly vortexed and heated for 5 min at 95 °C. Sample volumes corresponding to 4 µg, 2 µg and 1 µg of total protein, alongside defined masses (25 ng, 50 ng, 100 ng) of tubulin purified from ESCs, were loaded onto NuPAGE 4–12% Bis-Tris protein gels (Thermo) and mounted in gel running chambers filled with 1x NuPAGE SDS running buffer (Thermo). A replicate gel (with adjusted mass of tubulin controls of 100 ng, 200 ng and 400 ng) was loaded in parallel for downstream Coomassie staining of total protein for normalisation ^111^. Electrophoresis was for (1) 15 min at 50V and (2) 60 min at 150V.

The original gel was blotted onto a nitrocellulose membrane using 1x NuPAGE transfer buffer and a wet transfer protocol in an ice cold transfer chamber for 1 h at 40V. The membrane was blocked in a ready-to-use TBS blocking buffer (LI-COR) for 1h at RT. Staining with monoclonal anti-alpha-tubulin antibodies (DM1a (Sigma T-6199), 1:5,000) was carried out in TBS blocking buffer supplemented with 0.2% Tween-20 overnight at 4 °C. Next, the membranes were washed three times using TBS-T and then incubated with secondary antibodies (AlexaFluor(800 nm) anti-mouse IgG (Thermo), 1:5,000) for 1 h at RT protected from light. Finally, the membranes were washed again three times in TBS-T and finally stored in TBS without detergent. Bands were fluorescently detected on an Odyssey XF imager system (LI-COR) with an integration time of 2 min. The raw signals were quantified using *FIJI*’s rectangular selection tool. First, the sum signals of the tubulin standards were measured to generate a calibration curve. Finally, the sum signals of the lysate lanes were determined (using the rectangular selection tool) and then calibrated using the calibration curve.

For downstream normalisation, the replicate gel was stained using an InstantBlue Coomassie protein stain (Abcam) according to the manufacturer’s instructions. Coomassie can be used as a sensitive fluorescent in-gel protein stain (Butt et al. 2013). Protein was fluorescently detected in a ChemiDoc system (Bio-Rad) using 700 nm excitation light. The raw total protein reference image was de-noised in *FIJI* using a median filter (sigma = 3 pixels). The background was subtracted using the FIJI rolling ball algorithm at a radius of 50 pixels. For each of the lanes including the tubulin calibration lanes, the raw pixel sum was measured using the rectangular selection tool.

### Western blotting

Whole-cell RIPA lysates (see above) were diluted 1:4 with SDS sample buffer (4x), briefly vortexed and heated for 5 min at 95 °C. Sample volumes corresponding to 4 µg of total protein were loaded onto NuPAGE 4–12% Bis-Tris protein gels (Thermo) and mounted in gel running chambers filled with 1x NuPAGE SDS running buffer (Thermo). Electrophoresis was performed in two steps: (1) 15 min at 50V and (2) 60 min at 150V. The gel was blotted onto a PVDF membrane (methanol activated) using 1x NuPAGE transfer buffer and a wet transfer protocol in an ice cold transfer chamber for 1 h at 40V. The membranes were incubated for 1 h in blocking buffer (5% skimmed milk powder in 1x TBS-T buffer) at RT. For immunodetection, primary antibodies were added to the blocking buffer and incubated overnight at 4 °C. After three washing steps with 1x TBS-T, the membranes were incubated with the secondary antibodies (horseradish peroxidase-conjugated, 1:15,000 diluted in blocking buffer) for 1 h at RT. After another three washing steps with 1x TBS-T, the membranes were placed in a ChemiDoc gel documentation system (Bio-Rad) for chemiluminescent detection (Clarity Max ECL substrate kit, Bio-Rad). The raw images were exported and analysed in *FIJI*.

### Tubulin affinity purification

Functional tubulin heterodimers were affinity purified using TOG columns ^68,112^. Cultures of undifferentiated ESCs and early-differentiated ESCs were scaled up using 35-40 culturing dishes (150 mm). Cells were harvested by Accutase treatment at RT for 5 min. After adding an equal volume of 1x PBS, cell suspensions were distributed onto 50 mL conical tubes and centrifuged for 5 min at 300 x g at 4 °C. The supernatants were removed, the pellets were flash-frozen in liquid nitrogen and stored at -80 °C. On the day of purification, the pellets were unified into a single conical 50 mL tube and thawed on ice. The thawed cell suspension was taken up in an equal volume of ice cold 1x BRB80 (80 mM PIPES, 1 mM MgCl2, and 1 mM EGTA, pH 6.9) supplemented with benzonase, DTT, protease inhibitors, deacetylase inhibitor and PhosStop. Lysis was performed on ice using a dounce homogeniser (“B” pestle). The remaining purification steps were carried out as described in detail in Reusch et al. 2020 (starting at step 25).

### Intact protein analysis by liquid chromatography mass spectrometry

Intact protein masses were determined by liquid chromatography mass spectrometry (LC-MS) as described in ^113^. In brief, samples were analysed using the Ultimate 3000 liquid chromatography system connected to a Q Exactive HF mass spectrometer via the ion max source with HESI-II probe (Thermo). The proteins were desalted and concentrated by injection on a reversed-phase cartridge (MSPac DS-10, 2.1×10 mm, Thermo). Full MS spectra were acquired using the following parameters: intact protein mode on, mass range m/z 600–2500, resolution 15,000, AGC target 3×10^6^, Microscans 5, maximum injection time 200 ms. The software tool *UniDec* was used for data processing and deconvolution of protein masses ^114^. First, an averaged spectrum was generated from each measurement followed by spectral deconvolution using the default settings except for the following adjustments: charge range 38-65, Mass range 45-55 kDa, sample mass every 1 Da, peak FWHM 0.1.

### Osmotic challenge

ESCs were seeded at a density of 19,000 cells per cm^2^ onto L511-coated surfaces and initially incubated in FBS+LIF medium overnight. The next day, the growth medium was diluted down to 75% of its original concentration with ultrapure DNase/RNase-free distilled water (Invitrogen). For the isosmotic control, full growth medium was added. To allow spindles to recover, live-cell imaging or chemical fixation was performed after 2.5 hours post-treatment. Media osmolalities were measured using freezing point osmometry via the Osmomat 3000 basic (Gonotec) following the manufacturer’s instructions.

### Quantification of cellular mass density

Cellular mass density was measured using a correlative optical diffraction tomography (ODT) and fluorescence microscopy setup. ESCs were seeded onto L511-coated 35mm polymer bottom imaging dishes (Ibidi) at a density of 50,000 cells per cm^2^ and incubated for 4 h at 37 °C, 95% humidity and 5% CO_2_. Refractive index (RI) tomograms by ODT were obtained on the custom-built setup as described in ^115^. In brief, a coherent laser beam with a wavelength of 532 nm into two using a beam splitter (BS028, Thorlabs Inc.). While one beam was used as a reference beam, the sample was illuminated with the other using a 60x water-dipping objective lens (LUMPLFLN60XW, NA = 1.0, Olympus Life Science). Using a dual-axis galvanometer mirror (GVS012/M, Thorlabs Inc.), the sample was illuminated from 150 incident angles. After passing through the sample, the scattered light was gathered by a 60x objective (oil immersion, UplanSApo, NA =1.35, Olympus Life Science). At the image plane, the sample beam interfered with the reference beam to generate spatially modulated holograms. These were recorded by a charge-coupled device (CCD) camera (FL3-U3-13Y3M-C, FLIR Systems, Inc.). To generate 3D tomograms, the holograms were reconstructed using a custom written *MATLAB* script (github.com/OpticalDiffractionTomography). More information regarding tomogram reconstruction can be found in ^116^.

In *FIJI*, the cells were 3D segmented as described above, based on the RI intensities in the tomograms. The average RI value within the 3D cell mask was determined using the *FIJI 3D Suite* ^110^. For translating the raw RI value into mass density (in mg/mL), the following formula was applied:

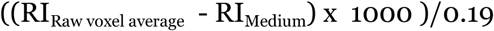

The refractive index of the medium (RI_Medium_) was determined for each experiment using an Abbe refractometer (Arcada ABBE-2WAJ). The cell dry mass is the product of the average cell mass density and the cell volume.

The ODT setup was built onto an IX71 microscope frame (Olympus Life Science) that was capable of epi-fluorescence imaging. The fluorescence images were acquired with the same objective used for ODT imaging and recorded onto a CMOS camera (FL-AB-20-BW 847 Tucsen Photonics Co., Ltd.).

## Data availability

All source data used in this study are deposited at Biostudies (TMP_17200187074432).

## Code availability

Source code, usage and installation guidelines for the implemented adaptive feedback microscopy pipeline is available in the following repository: https://git.embl.de/grp-almf/automictools-zenblue-spindle-kletter. Analytical codes used for this study are deposited under https://github.com/TobiasKletter/Scaling.

## Acknowledgements

The authors thank all past and current members of the Zaburdaev and Reber labs. We thank the AMBIO (Charité, Berlin) and the Advanced Light Microscopy Facility (ALMF) at the European Molecular Biology Laboratory (EMBL) and Zeiss for imaging support. For mass spectrometry, we would like to acknowledge the assistance of the Core Facility BioSupraMol supported by the Deutsche Forschungsgemeinschaft (DFG). We would like to thank Professor Dr. Jochen Guck and Dr. Kyoohyun Kim (Max Planck Institute for the Science of Light) for their support in the development of the correlative ODT-fluorescence setup and analysis pipeline. We thank Elena Taverna and Felipe Mora-Bermúdez for help and advice on neural differentiation and Christian Schlieker, Marcus Taylor, Valentina Štimac, and Manuel Mendoza for critical comments on the manuscript.

## Author contributions

T.K. and S.R. conceived and designed the study. T.K. performed all experiments including data and image analysis. S.R. contribute to method development. A.B. supported ODT setup development and quantitative phase imaging. B.N. supported long-term live cell imaging and RNAi experiments. A.H. designed the Adaptive Microscopy Feedback pipeline and supported these experiments. B.K. performed and interpreted the mass-spectrometric analyses. O.M. and V.Z. conceptualised and wrote the theory part. T.K. and S.R. wrote the manuscript with input from all authors.

## Funding

The authors acknowledge funding from the Max Planck Society (to O.M. and S.R.) and from the Horizon 2020 Framework Programme of the European Union (iNEXT grant 653706, project PID 3503 to S. R.).

## Competing interests

The authors declare no competing interests.

## Notes

### Competing Interest Statement

The authors have declared no competing interest.

## References

1. Thompson, D. W. On Growth and Form. (Cambridge University Press, 1992).

2. Valdez, V. A., Neahring, L., Petry, S. & Dumont, S. Mechanisms underlying spindle assembly and robustness. Nat. Rev. Mol. Cell Biol. 24, 523–542 (2023).

3. Wühr, M. et al. Evidence for an Upper Limit to Mitotic Spindle Length. Curr. Biol. 18, 1256–1261 (2008).

4. Greenan, G. et al. Centrosome Size Sets Mitotic Spindle Length in Caenorhabditis elegans Embryos. Curr. Biol. 20, 353–358 (2010).

5. Courtois, A., Schuh, M., Ellenberg, J. & Hiiragi, T. The transition from meiotic to mitotic spindle assembly is gradual during early mammalian development. J. Cell Biol. 198, 357–370 (2012).

6. Wilbur, J. D. & Heald, R. Mitotic spindle scaling during Xenopus development by kif2a and importin α. Elife 2013, 2–4 (2013).

7. Farhadifar, R. et al. Scaling, selection, and evolutionary dynamics of the mitotic spindle. Curr. Biol. 25, 732–740 (2015).

8. Milunovic-Jevtic, A., Jevtic, P., Levy, D. L. & Gatlin, J. C. In vivo mitotic spindle scaling can be modulated by changing the levels of a single protein: the microtubule polymerase XMAP215. Mol. Biol. Cell 29, 1311–1317 (2018).

9. Rieckhoff, E. M. et al. Spindle Scaling Is Governed by Cell Boundary Regulation of Microtubule Nucleation. Curr. Biol. 30, 4973–4983.e10 (2020).

10. Kiyomitsu, A. et al. Ran-GTP assembles a specialized spindle structure for accurate chromosome segregation in medaka early embryos. Nat. Commun. 15, 981 (2024).

11. Kletter, T., Biswas, A. & Reber, S. Engineering metaphase spindles: Construction site and building blocks. Curr. Opin. Cell Biol. 79, 102143 (2022).

12. Decker, F., Oriola, D., Dalton, B. & Brugués, J. Autocatalytic microtubule nucleation determines the size and mass of Xenopus laevis egg extract spindles. Elife 7, (2018).

13. Lacroix, B. et al. Microtubule Dynamics Scale with Cell Size to Set Spindle Length and Assembly Timing. Dev. Cell 45, 496–511.e6 (2018).

14. Hirst, W. G., Biswas, A., Mahalingan, K. K. & Reber, S. Differences in Intrinsic Tubulin Dynamic Properties Contribute to Spindle Length Control in Xenopus Species. Curr. Biol. 30, 2184–2190.e5 (2020).

15. Loughlin, R., Wilbur, J. D., McNally, F. J., Nédélec, F. J. & Heald, R. Katanin contributes to interspecies spindle length scaling in xenopus. Cell 147, 1397–1407 (2011).

16. Helmke, K. J. & Heald, R. TPX2 levels modulate meiotic spindle size and architecture in Xenopus egg extracts. J. Cell Biol. 206, 385–393 (2014).

17. Miller, K. E., Session, A. M. & Heald, R. Kif2a Scales Meiotic Spindle Size in Hymenochirus boettgeri. Curr. Biol. 29, 3720–3727.e5 (2019).

18. Good, M. C., Vahey, M. D., Skandarajah, A., Fletcher, D. A. & Heald, R. Cytoplasmic volume modulates spindle size during embryogenesis. Science 342, 856–860 (2013).

19. Hazel, J. et al. Changes in cytoplasmic volume are sufficient to drive spindle scaling. Science 342, 853–856 (2013).

20. Brownlee, C. & Heald, R. Importin a Partitioning to the Plasma Membrane Regulates Intracellular Scaling. Cell 176, 805–815.e8 (2019).

21. Mora-Bermúdez, F., Matsuzaki, F. & Huttner, W. B. Specific polar subpopulations of astral microtubules control spindle orientation and symmetric neural stem cell division. Elife 3, 1–31 (2014).

22. Vargas-Hurtado, D. et al. Differences in Mitotic Spindle Architecture in Mammalian Neural Stem Cells Influence Mitotic Accuracy during Brain Development. Curr. Biol. 29, 2993–3005.e9 (2019).

23. Molines, A. T. et al. Physical properties of the cytoplasm modulate the rates of microtubule polymerization and depolymerization. Dev. Cell 57, 466–479.e6 (2022).

24. Shimamoto, Y., Maeda, Y. T., Ishiwata, S.‘ichi, Libchaber, A. J. & Kapoor, T. M. Insights into the micromechanical properties of the metaphase spindle. Cell 145, 1062–1074 (2011).

25. Bai, L. & Mitchison, T. J. Spring-like behavior of cytoplasm holds the mitotic spindle in place. Proceedings of the National Academy of Sciences of the United States of America vol. 119 e2203036119 (2022).

26. Xie, J. et al. Contribution of cytoplasm viscoelastic properties to mitotic spindle positioning. Proc. Natl. Acad. Sci. U. S. A. 119, (2022).

27. Mchedlishvili, N., Matthews, H. K., Corrigan, A. & Baum, B. Two-step interphase microtubule disassembly aids spindle morphogenesis. BMC Biol. 16, 14 (2018).

28. Schweizer, N., Pawar, N., Weiss, M. & Maiato, H. An organelle-exclusion envelope assists mitosis and underlies distinct molecular crowding in the spindle region. J. Cell Biol. 210, 695–704 (2015).

29. Biswas, A., Kim, K., Cojoc, G., Guck, J. & Reber, S. The Xenopus spindle is as dense as the surrounding cytoplasm. Dev. Cell 56, 967–975.e5 (2021).

30. Woodruff, J. B. et al. The Centrosome Is a Selective Condensate that Nucleates Microtubules by Concentrating Tubulin. Cell 169, 1066–1077.e10 (2017).

31. Baumgart, J. et al. Soluble tubulin is significantly enriched at mitotic centrosomes. J. Cell Biol. 218, 3977–3985 (2019).

32. Hurst, S., Vos, B. E., Brandt, M. & Betz, T. Intracellular softening and increased viscoelastic fluidity during division. Nat. Phys. 17, 1270–1276 (2021).

33. Carlini, L., Brittingham, G. P., Holt, L. J. & Kapoor, T. M. Microtubules Enhance Mesoscale Effective Diffusivity in the Crowded Metaphase Cytoplasm. Dev. Cell 54, 574–582.e4 (2020).

34. Maric, D., Maric, I. & Barker, J. L. Buoyant Density Gradient Fractionation and Flow Cytometric Analysis of Embryonic Rat Cortical Neurons and Progenitor Cells. Methods 16, 247–259 (1998).

35. Jackson, C. W., Steward, S. A., Brown, L. K. & Look, A. T. Inverse relationship between megakaryocyte buoyant density and maturity. Br. J. Haematol. 64, 33–43 (1986).

36. Sasai, Y., Hachisuka, H., Mori, O. & Nomura, H. Separation of keratinocytes by density gradient centrifugation for DNA cytofluorometry. Histochemistry 80, 133–136 (1984).

37. Chalut, K. J., Ekpenyong, A. E., Clegg, W. L., Melhuish, I. C. & Guck, J. Quantifying cellular differentiation by physical phenotype using digital holographic microscopy. Integr. Biol. 4, 280 (2012).

38. Cooper, K. L. et al. Multiple phases of chondrocyte enlargement underlie differences in skeletal proportions. Nature 495, 375–378 (2013).

39. Neurohr, G. E. & Amon, A. Relevance and Regulation of Cell Density. Trends Cell Biol. 30, 213–225 (2020).

40. Bergert, M. et al. Cell Surface Mechanics Gate Embryonic Stem Cell Differentiation. Cell Stem Cell 28, 209–216.e4 (2021).

41. Woods, C. G. & Basto, R. Microcephaly. Curr. Biol. 24, R1109–R1111 (2014).

42. Nano, M. & Basto, R. The Janus soul of centrosomes: a paradoxical role in disease? Chromosome Res. 24, 127–144 (2016).

43. Viais, R. et al. Augmin deficiency in neural stem cells causes p53-dependent apoptosis and aborts brain development. Elife 10, (2021).

44. Zheng, X. et al. Conserved TCP domain of Sas-4/CPAP is essential for pericentriolar material tethering during centrosome biogenesis. Proceedings of the National Academy of Sciences 111, (2014).

45. Ramani, A. et al. Plk1/Polo Phosphorylates Sas-4 at the Onset of Mitosis for an Efficient Recruitment of Pericentriolar Material to Centrosomes. Cell Rep. 25, 3618–3630.e6 (2018).

46. Varadarajan, R. & Rusan, N. M. Bridging centrioles and PCM in proper space and time. Essays Biochem. 62, 793–801 (2018).

47. Ying, Q. L., Stavridis, M., Griffiths, D., Li, M. & Smith, A. Conversion of embryonic stem cells into neuroectodermal precursors in adherent monoculture. Nat. Biotechnol. 21, 183–186 (2003).

48. Wongpaiboonwattana, W. & Stavridis, M. P. Neural differentiation of mouse embryonic stem cells in serum-free monolayer culture. J. Vis. Exp. 2015, 45–49 (2015).

49. Taverna, E., Götz, M. & Huttner, W. B. The Cell Biology of Neurogenesis: Toward an Understanding of the Development and Evolution of the Neocortex. Annu. Rev. Cell Dev. Biol. 30, 465–502 (2014).

50. Miotto, M. et al. Collective behavior and self-organization in neural rosette morphogenesis. Frontiers in Cell and Developmental Biology 11, (2023).

51. Kletter, T. et al. Volumetric morphometry reveals spindle width as the best predictor of mammalian spindle scaling. J. Cell Biol. 221, (2021).

52. Berg, S. et al. Ilastik: Interactive Machine Learning for (Bio)Image Analysis. Nat. Methods 16, 1226–1232 (2019).

53. Reber, S. B. et al. XMAP215 activity sets spindle length by controlling the total mass of spindle microtubules. Nat. Cell Biol. 15, 1116–1122 (2013).

54. Brugués, J. & Needleman, D. Physical basis of spindle self-organization. Proc. Natl. Acad. Sci. U. S. A. 111, 18496–18500 (2014).

55. Gruss, O. J. et al. Ran induces spindle assembly by reversing the inhibitory effect of importin α on TPX2 activity. Cell 104, 83–93 (2001).

56. Roostalu, J., Cade, N. I. & Surrey, T. Complementary activities of TPX2 and chTOG constitute an efficient importin-regulated microtubule nucleation module. Nat. Cell Biol. 17, 1422–1434 (2015).

57. Alfaro-Aco, R., Thawani, A. & Petry, S. Biochemical reconstitution of branching microtubule nucleation. Elife 9, 1–16 (2020).

58. Srayko, M., O’Toole, E. T., Hyman, A. A. & Müller-Reichert, T. Katanin Disrupts the Microtubule Lattice and Increases Polymer Number in C. elegans Meiosis. Curr. Biol. 16, 1944–1949 (2006).

59. McNally, K., Audhya, A., Oegema, K. & McNally, F. J. Katanin controls mitotic and meiotic spindle length. J. Cell Biol. 175, 881–891 (2006).

60. Loughlin, R., Heald, R. & Nédélec, F. A computational model predicts Xenopus meiotic spindle organization. J. Cell Biol. 191, 1239–1249 (2010).

61. Guerreiro, A. et al. Wdr62 localizes katanin at spindle poles to ensure synchronous chromosome segregation. J. Cell Biol. 220, (2021).

62. Piehl, M., Tulu, U. S., Wadsworth, P. & Cassimeris, L. Centrosome maturation: Measurement of microtubule nucleation throughout the cell cycle by using GFP-tagged EB1. Proceedings of the National Academy of Sciences 101, 1584–1588 (2004).

63. Woodruff, J. B., Wueseke, O. & Hyman, A. A. Pericentriolar material structure and dynamics. Philos. Trans. R. Soc. Lond. B Biol. Sci. 369, 20130459 (2014).

64. Gomes Pereira, S., Dias Louro, M. A. & Bettencourt-Dias, M. Biophysical and Quantitative Principles of Centrosome Biogenesis and Structure. Annu. Rev. Cell Dev. Biol. 37, 43–63 (2021).

65. Decker, M. et al. Limiting amounts of centrosome material set centrosome size in C. elegans embryos. Curr. Biol. 21, 1259–1267 (2011).

66. Rathbun, L. I. et al. PLK1- and PLK4-Mediated Asymmetric Mitotic Centrosome Size and Positioning in the Early Zebrafish Embryo. Curr. Biol. 30, 4519–4527.e3 (2020).

67. Liu, P. et al. Insights into the assembly and activation of the microtubule nucleator γ-TuRC. Nature 578, 467–471 (2020).

68. Reusch, S., Biswas, A., Hirst, W. G. & Reber, S. Affinity Purification of Label-free Tubulins from Xenopus Egg Extracts. STAR Protocols 1, 100151 (2020).

69. Kim, K. & Guck, J. The Relative Densities of Cytoplasm and Nuclear Compartments Are Robust against Strong Perturbation. Biophys. J. 119, 1946–1957 (2020).

70. Gopalakrishnan, J. et al. Tubulin nucleotide status controls Sas-4-dependent pericentriolar material recruitment. Nat. Cell Biol. 14, 865–873 (2012).

71. Mariappan, A. et al. Inhibition of CPAP –tubulin interaction prevents proliferation of centrosome-amplified cancer cells. EMBO J. 38, (2019).

72. Crowder, M. E. et al. A comparative analysis of spindle morphometrics across metazoans. Curr. Biol. 25, 1542–1550 (2015).

73. Engler, A. J., Sen, S., Sweeney, H. L. & Discher, D. E. Matrix elasticity directs stem cell lineage specification. Cell 126, 677–689 (2006).

74. Guo, M. et al. Cell volume change through water efflux impacts cell stiffness and stem cell fate. Proc. Natl. Acad. Sci. U. S. A. 114, E8618–E8627 (2017).

75. Rafelski, S. M. & Theriot, J. A. Establishing a conceptual framework for holistic cell states and state transitions. Cell 187, 2633–2651 (2024).

76. Newport, J. & Kirschner, M. A major developmental transition in early xenopus embryos: II. control of the onset of transcription. Cell 30, 687–696 (1982).

77. Peshkin, L. et al. On the Relationship of Protein and mRNA Dynamics in Vertebrate Embryonic Development. Dev. Cell 35, 383–394 (2015).

78. Cuomo, A. S. E. et al. Single-cell RNA-sequencing of differentiating iPS cells reveals dynamic genetic effects on gene expression. Nat. Commun. 11, 810 (2020).

79. Ochocka, N. et al. Single-cell RNA sequencing reveals functional heterogeneity of glioma-associated brain macrophages. Nat. Commun. 12, 1151 (2021).

80. Mulas, C., Chaigne, A., Smith, A. & Chalut, K. J. Cell state transitions: definitions and challenges. Development 148, (2021).

81. Al-Dosari, M. S., Shaheen, R., Colak, D. & Alkuraya, F. S. Novel CENPJ mutation causes Seckel syndrome. J. Med. Genet. 47, 411–414 (2010).

82. Kitagawa, D. et al. Spindle positioning in human cells relies on proper centriole formation and on the microcephaly proteins CPAP and STIL. J. Cell Sci. 124, 3884–3893 (2011).

83. An, H.-L., Kuo, H.-C. & Tang, T. K. Modeling Human Primary Microcephaly With hiPSC-Derived Brain Organoids Carrying CPAP-E1235V Disease-Associated Mutant Protein. Frontiers in Cell and Developmental Biology 10, (2022).

84. David, A. F. et al. Augmin accumulation on long-lived microtubules drives amplification and kinetochore-directed growth. J. Cell Biol. 218, 2150–2168 (2019).

85. Khodjakov, A. & Rieder, C. L. Centrosomes enhance the fidelity of cytokinesis in vertebrates and are required for cell cycle progression. J. Cell Biol. 153, 237–242 (2001).

86. Nichols, J. et al. Formation of Pluripotent Stem Cells in the Mammalian Embryo Depends on the POU Transcription Factor Oct4. Cell 95, 379–391 (1998).

87. Arai, Y. et al. Neural stem and progenitor cells shorten S-phase on commitment to neuron production. Nat. Commun. 2, (2011).

88. Waisman, A. et al. Cell cycle dynamics of mouse embryonic stem cells in the ground state and during transition to formative pluripotency. Sci. Rep. 9, (2019).

89. Zhai, Y., Kronebusch, P. J., Simon, P. M. & Borisy, G. G. Microtubule dynamics at the G2/M transition: abrupt breakdown of cytoplasmic microtubules at nuclear envelope breakdown and implications for spindle morphogenesis. J. Cell Biol. 135, 201–214 (1996).

90. Brugués, J., Nuzzo, V., Mazur, E. & Needleman, D. J. Nucleation and transport organize microtubules in metaphase spindles. Cell 149, 554–564 (2012).

91. Needleman, D. J. et al. Fast microtubule dynamics in meiotic spindles measured by single molecule imaging: evidence that the spindle environment does not stabilize microtubules. Mol. Biol. Cell 21, 323–333 (2010).

92. Goodson, H. V. & Jonasson, E. M. Microtubules and Microtubule-Associated Proteins. Cold Spring Harb. Perspect. Biol. 10, (2018).

93. Sharma, A. et al. Centriolar CPAP/SAS-4 Imparts Slow Processive Microtubule Growth. Dev. Cell 37, 362–376 (2016).

94. Ton, C. C. T. et al. Positional cloning and characterization of a paired box- and homeobox-containing gene from the aniridia region. Cell 67, 1059–1074 (1991).

95. Frederiksen, K. & McKay, R. D. Proliferation and differentiation of rat neuroepithelial precursor cells in vivo. J. Neurosci. 8, 1144–1151 (1988).

96. Guillemot, F. et al. Mammalian achaete-scute homolog 1 is required for the early development of olfactory and autonomic neurons. Cell 75, 463–476 (1993).

97. Aaku-Saraste, E., Hellwig, A. & Huttner, W. B. Loss of Occludin and Functional Tight Junctions, but Not ZO-1, during Neural Tube Closure—Remodeling of the Neuroepithelium Prior to Neurogenesis. Dev. Biol. 180, 664–679 (1996).

98. Arnold, S. J. et al. The T-box transcription factor Eomes/Tbr2 regulates neurogenesis in the cortical subventricular zone. Genes Dev. 22, 2479–2484 (2008).

99. Camargo Ortega, G. et al. The centrosome protein AKNA regulates neurogenesis via microtubule organization. Nature 567, 113–117 (2019).

100. Reback, J. et al. pandas-dev/pandas: Pandas 1.1.3. (2020) doi:10.5281/ZENODO.4067057.

101. Harris, C. R. et al. Array programming with NumPy. Nature 585, 357–362 (2020).

102. Virtanen, P. et al. SciPy 1.0: fundamental algorithms for scientific computing in Python. Nat. Methods 17, 261–272 (2020).

103. Seabold, S. & Perktold, J. Statsmodels: Econometric and Statistical Modeling with Python. http://statsmodels.sourceforge.net/ (2010).

104. Pedregosa, F. et al. Scikit-Learn: Machine Learning in Python. vol. 12 2825–2830 http://scikit-learn.sourceforge.net. (2011).

105. Poser, I. et al. BAC TransgeneOmics: a high-throughput method for exploration of protein function in mammals. Nat. Methods 5, 409–415 (2008).

106. Schindelin, J. et al. Fiji: An open-source platform for biological-image analysis. Nat. Methods 9, 676–682 (2012).

107. Thomas, L. S. V. & Gehrig, J. Multi-template matching: a versatile tool for object-localization in microscopy images. BMC Bioinformatics 21, 44 (2020).

108. Tinevez, J.-Y. et al. TrackMate: An open and extensible platform for single-particle tracking. Methods 115, 80–90 (2017).

109. Legland, D., Arganda-Carreras, I. & Andrey, P. MorphoLibJ: Integrated library and plugins for mathematical morphology with ImageJ. Bioinformatics 32, 3532–3534 (2016).

110. Ollion, J., Cochennec, J., Loll, F., Escudé, C. & Boudier, T. TANGO: A generic tool for high-throughput 3D image analysis for studying nuclear organization. Bioinformatics 29, 1840–1841 (2013).

111. Eaton, S. L. et al. Total Protein Analysis as a Reliable Loading Control for Quantitative Fluorescent Western Blotting. PLoS One 8, e72457 (2013).

112. Widlund, P. O. et al. One-step purification of assembly-competent tubulin from diverse eukaryotic sources. Mol. Biol. Cell 23, 4393–4401 (2012).

113. Hirst, W. G. et al. Purification of functional Plasmodium falciparum tubulin allows for the identification of parasite-specific microtubule inhibitors. Curr. Biol. 32, 919–926.e6 (2022).

114. Marty, M. T. et al. Bayesian deconvolution of mass and ion mobility spectra: from binary interactions to polydisperse ensembles. Anal. Chem. 87, 4370–4376 (2015).

115. Biswas, A. et al. Conserved nucleocytoplasmic density homeostasis drives cellular organization across eukaryotes. bioRxiv 2023.09.05.556409 (2023) doi:10.1101/2023.09.05.556409.

116. Kim, K. et al. High-resolution three-dimensional imaging of red blood cells parasitized by Plasmodium falciparum and in situ hemozoin crystals using optical diffraction tomography. J. Biomed. Opt. 19, 011005 (2014).

